# Computation noise promotes cognitive resilience to adverse conditions during decision-making

**DOI:** 10.1101/2020.06.10.145300

**Authors:** Charles Findling, Valentin Wyart

**Affiliations:** Laboratoire de Neurosciences Cognitives et Computationnelles, Institut National de la Santé et de la Recherche Médicale, Paris, France; Département d’Études Cognitives, École Normale Supérieure, Université PSL, Paris, France

## Abstract

Random noise in information processing systems is widely seen as detrimental to function. But despite the large trial-to-trial variability of neural activity and behavior, humans and other animals show a remarkable adaptability to unexpected adverse events occurring during task execution. This cognitive ability, described as constitutive of general intelligence, is missing from current artificial intelligence (AI) systems which feature exact (noise-free) computations. Here we show that implementing computation noise in recurrent neural networks boosts their cognitive resilience to a variety of adverse conditions entirely unseen during training, in a way that resembles human and animal cognition. In contrast to artificial agents with exact computations, noisy agents exhibit hallmarks of Bayesian inference acquired in a ‘zero-shot’ fashion – without prior experience with conditions that require these computations for maximizing rewards. We further demonstrate that these cognitive benefits result from free-standing regularization of activity patterns in noisy neural networks. Together, these findings suggest that intelligence may ride on computation noise to promote near-optimal decision-making in adverse conditions without any engineered cognitive sophistication.

## Introduction

Extracting signal from noise is seen as a core feature of efficient information processing systems, from gravitational-wave detectors to neural networks. In this context, noise is usually defined as irrelevant input that should be filtered out to improve signal detection. In recent years, artificial neural networks have reached high expertise when it comes to extracting signal from input noise in widely different situations^1^. After training, which typically requires hundreds of thousands of iterations, deep neural networks (DNNs) are able to reach human-level and even superhuman performance in a large variety of signal detection tasks, including the visual categorization of challenging visual patterns^2^ and natural images^3^.

However, these large networks are ‘paper tigers’ when it comes to adapting to novel conditions not encountered during training – i.e., beyond out-of-sample instances of the condition used for training^4,5^. As a prime example, artificial agents trained to maximize rewards by identifying predictive signals in their environment often fail to adapt to novel environments, even ones conceptually very similar to the ones seen during training, and to adverse events occurring in the same environment^6,7^. A deep network trained on one level of a video game usually fares surprisingly badly on other – let alone harder – levels of the same game^8^. These ‘generalization failures’ are seen as one of the main challenges for achieving artificial general intelligence (AGI). Indeed, it is the ability to acquire new skills from previously acquired skills that defines general intelligence according to current definitions, not the level of expertise at any given task^9,10^ – whether it is categorizing dogs vs. cats, playing Atari games^11^ or mastering the game of Go^12^.

In stark contrast to artificial agents, biological agents – humans in particular – show much higher cognitive resilience to a variety of unpredictable adverse events, or ‘shocks’, occurring in their environments. In controlled laboratory conditions, adverse conditions faced by human agents include the presentation of conflicting pieces of information, but also sudden changes in environment statistics during learning and decision-making. Influential theories^13,14^ postulate that humans developed purposeful heuristics to respond efficiently to these different shocks – a ‘bag of tricks’ that deep networks lack unless they are exposed to all possible adverse events during training.

Here we propose a radically different account of the high cognitive resilience of human agents to adverse conditions, arising from a core property of biological neural networks not shared with their artificial counterparts. Beyond input noise, biological neural networks process and respond to input with a large internally generated variability^15^ that is absent from artificial neural networks whose units exhibit deterministic input-state-output relations after training. This ‘computation noise’ has a wide-ranging impact on human cognition and behavior^16^ – from fluctuations in the perception of weak sensory stimuli^17^ to ‘exploratory’ behavior during reward-guided decision-making^18–20^. Existing research has typically considered this internal noise as a hard constraint on neural information processing systems, that the brain has evolved to cope with using efficient coding strategies^21–24^.

Instead, we hypothesized that the large computation noise observed in neural systems and the resulting behavior may serve a specific function: to promote the high cognitive resilience characteristic of human agents by providing online, free-standing regularization in neural networks. Instead of engineering cognitive resilience to adverse conditions through sophistication of neural networks, we tested the adaptive function of an apparent limitation of human cognition – its limited computational precision – by implementing it in artificial neural networks.

In practice, we compared recurrent neural networks (RNNs) – a canonical type of artificial neural networks used as model of associative cortical circuits^25–28^ (e.g., in parietal and prefrontal regions) – trained and tested with noisy vs. exact computations (Fig. 1), in two widely used tasks for studying higher cognition in humans and other animals^29,30^. The noisy and exact agents were trained using reinforcement learning (RL), and the exact agents included an explicit action entropy regularization term in their objective function to encourage exploration during training^31,32^. We found that computation noise confers several non-trivial cognitive advantages to artificial neural networks, including zero-shot (training-free) adaptability to various types of adverse conditions never encountered during training.

**Fig. 1.**
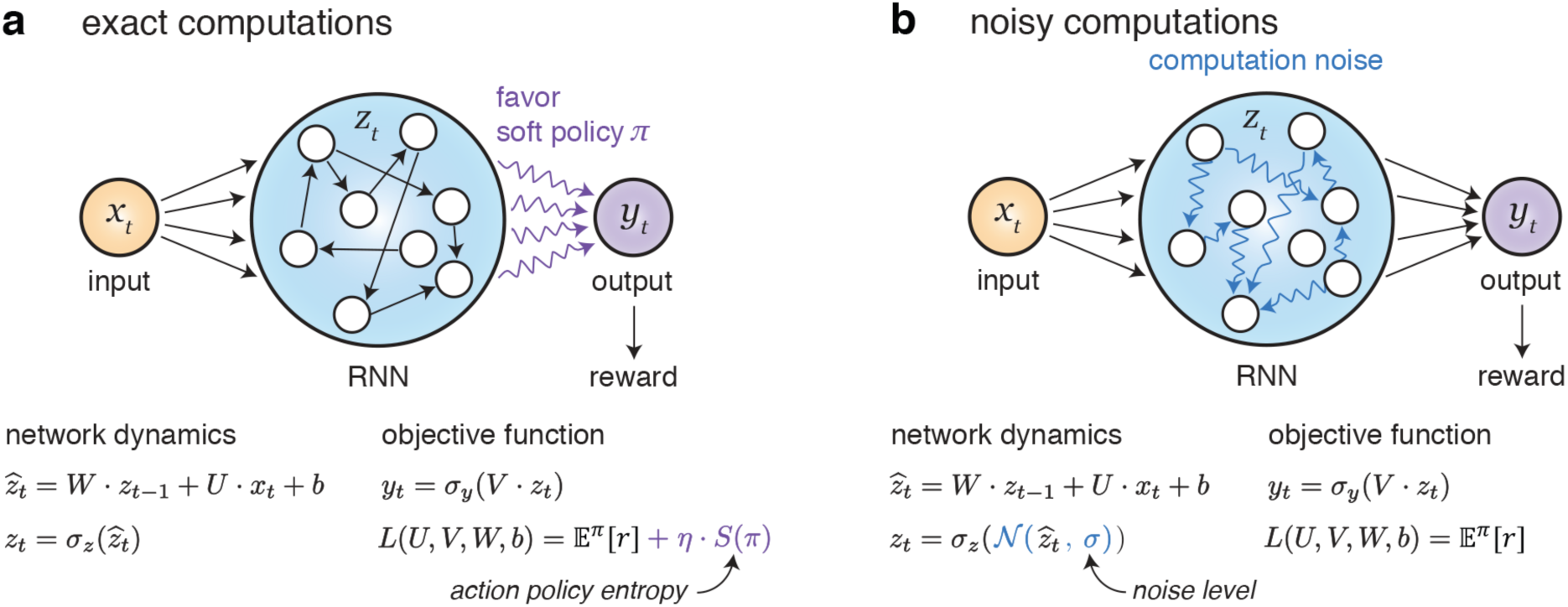
Description of the recurrent neural networks. The decision-making RNN is fed with input *x*_*t*_, which is combined with the previous recurrent activity *z*_*t*−1_ and passed through a non-linear activation function *σ*_*Z*_(·) to obtain the updated recurrent activity *z*_*t*_. The output (action policy) *y*_*t*_ of the RNN is obtained from *z*_*t*_, and passed through a softmax function *σ*_*Y*_(·) to choose an action. **a**, RNN with exact (noise-free) computations. The objective function of the network maximizes expected rewards, with an optional action entropy regularization term *S*(*π*) used to encourage exploration. **b**, RNN with noisy computations. The recurrent network updates are now corrupted by zero-mean normally distributed noise of standard deviation *σ*. The objective function of the network maximizes expected rewards, without any explicit regularization term.

## Results

### Zero-shot learning of the weather prediction task in noisy agents

The first task we considered is the ‘weather prediction’ task^33–35^, which typically proceeds in two successive stages (Fig. 2a). In the first, ‘associative learning’ stage, the agent learns the probabilistic associations between a set of symbols (cues) and rewarded actions. On each trial, the agent is presented with one of the cues for a period of time, after which the agent has to choose between two actions (here, a saccade toward the red target vs. a saccade toward the green target). The agent receives a positive or negative reward as a function of the association between the presented cue and the chosen action (e.g., the star is associated with a 0.7 reward probability if the red target is chosen, and a 0.3 reward probability if the green target is chosen). After the agent has learnt the reward probabilities associated with each cue through repeated trials, the same agent is presented in the ‘weather prediction’ stage with sequences of different cues. After each sequence, the agent has to choose between the same two actions the one most likely to be associated with reward given the presented cues. In this second stage, each cue taken in isolation is associated with the same reward probabilities as in the first stage, but reward probabilities have to be combined across presented cues to identify the best action associated with the sequence of cues. The difficulty of this ‘weather prediction’ stage can be manipulated by varying the fraction of conflicting cues (i.e., cues associated with a different action than the sequence as a whole; Fig. 2b).

**Fig. 2.**
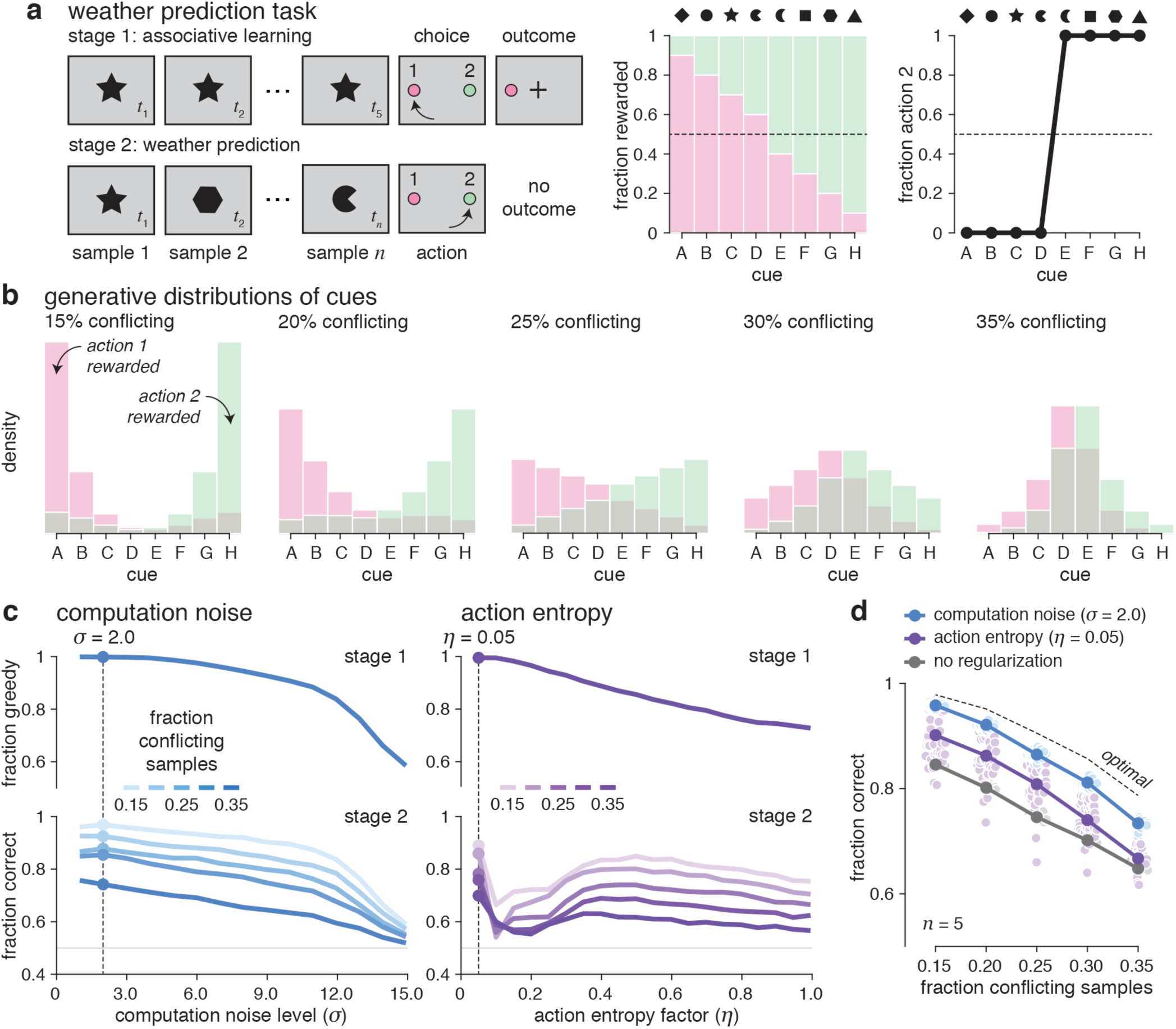
Weather prediction task. **a**, Description of the experimental paradigm. Left: in stage 1, the agent is presented with 5 samples of the same symbol (cue), then chooses between two actions and obtains a positive or negative outcome which depends on the probabilistic association between the presented cue and the chosen action. In stage 2, the same agent is now presented with sequences consisting of samples of different cues, then chooses between the same two actions the one most likely to be associated with reward given the sequence as a whole. Middle: fraction of trials for which each action is rewarded for each of the 8 cues. Right: fraction of trials for which action 2 is chosen in response to each cue after training in stage 1. **b**, Generative distributions of cues in stage 2, for trials in which action 1 is rewarded (red) and trials in which action 2 is rewarded (green). Conflicting cues are defined as cues whose associated action do not match the rewarded action of the sequence as a whole. We considered 5 levels of task difficulty in stage 2, corresponding to increasing fractions of conflicting cues in a sequence (from left to right). **c**, Fractions of ‘greedy’ decisions in stage 1 (top) and fraction of correct decisions in stage 2 for *n* = 5 samples (bottom) for noisy RNNs (left) and exact RNNs with action entropy regularization (right). As for all subsequent analyses, results were obtained by training 50 agents per computation noise level and per action entropy factor, and averaging their simulated behavior after training. Noisy RNNs performed substantially better than exact RNNs across a wide range of computation noise levels in stage 2. **d**, Fraction of correct decisions in stage 2 (*n* = 5 samples) as a function of task difficulty for noisy RNNs (blue), exact RNNs with action entropy regularization (violet) and non-regularized RNNs (gray). The black dashed line indicates the accuracy of the Bayes-optimal decision-maker. The small dots in the background indicate the accuracy of individual RNNs (50 per network type).

We trained RNNs featuring either noisy or exact computations to learn probabilistic associations in stage 1 through reinforcement learning (see Methods). In stage 1, we used a fixed (and arbitrary) presentation time of 5 samples before probing the RNNs for an action (Fig. 2a, top). We then tested the ability of these artificial agents to maximize reward not only in the task used during training (stage 1), but also in stage 2 – an adverse condition unseen during training. Stage 2 constitutes an adverse condition for agents trained in stage 1, because of the different and often conflicting cues presented at successive time samples of the same trial (Fig. 2a, bottom). We varied the level of computation noise (fixed across training and testing) in the noisy RNNs, and the amount of explicit action entropy regularization in the exact RNNs (see Methods). Computation noise was implemented by corrupting the updates of each unit in the network with i.i.d. normally distributed noise of standard deviation *σ*.

Although both types of agents performed optimally in stage 1 after training for moderate levels of computation noise and action entropy regularization (Fig. 2c, top), noisy RNNs performed substantially better than their exact counterparts across a wide range of computation noise levels in stage 2 (Fig. 2c, bottom). Extracting the computation noise level and the action entropy factor which maximized accuracy in stage 2 while retaining optimal performance in stage 1 revealed that noisy RNNs strongly outperformed exact RNNs across sequences of increasing difficulty (Fig. 2d), reaching near-optimal accuracy in a task condition entirely unseen during training. Optimality in stage 2 is defined as the greedy selection of the action associated with the highest posterior probability of reward. This posterior probability is obtained through Bayesian inference, by combining the reward probabilities associated with each cue in the sequence.

### Hallmarks of Bayesian inference in noisy agents

A hallmark of Bayesian inference in the weather prediction task is that accuracy should increase monotonically with more samples in the sequence. To test whether noisy RNNs outperformed exact RNNs by performing computations which resemble Bayesian inference, we presented artificial agents with sequences of cues of varying lengths, from 2 to 16 samples (Fig. 3a). As predicted, we found that the accuracy of noisy RNNs in stage 2 grew parametrically with sequence length, a qualitative pattern not found for exact RNNs (Fig. 3a, left). Explicit action entropy regularization during training even led to a clear case of ‘overfitting’, in the sense that exact RNNs trained with a presentation time of 5 samples were unable to make accurate decisions for other presentation times, even in stage 1 (Fig. 3a, middle). The accuracy of non-regularized RNNs was independent of the number of samples – something inconsistent with Bayesian inference (Fig. 3a, right).

**Fig. 3.**
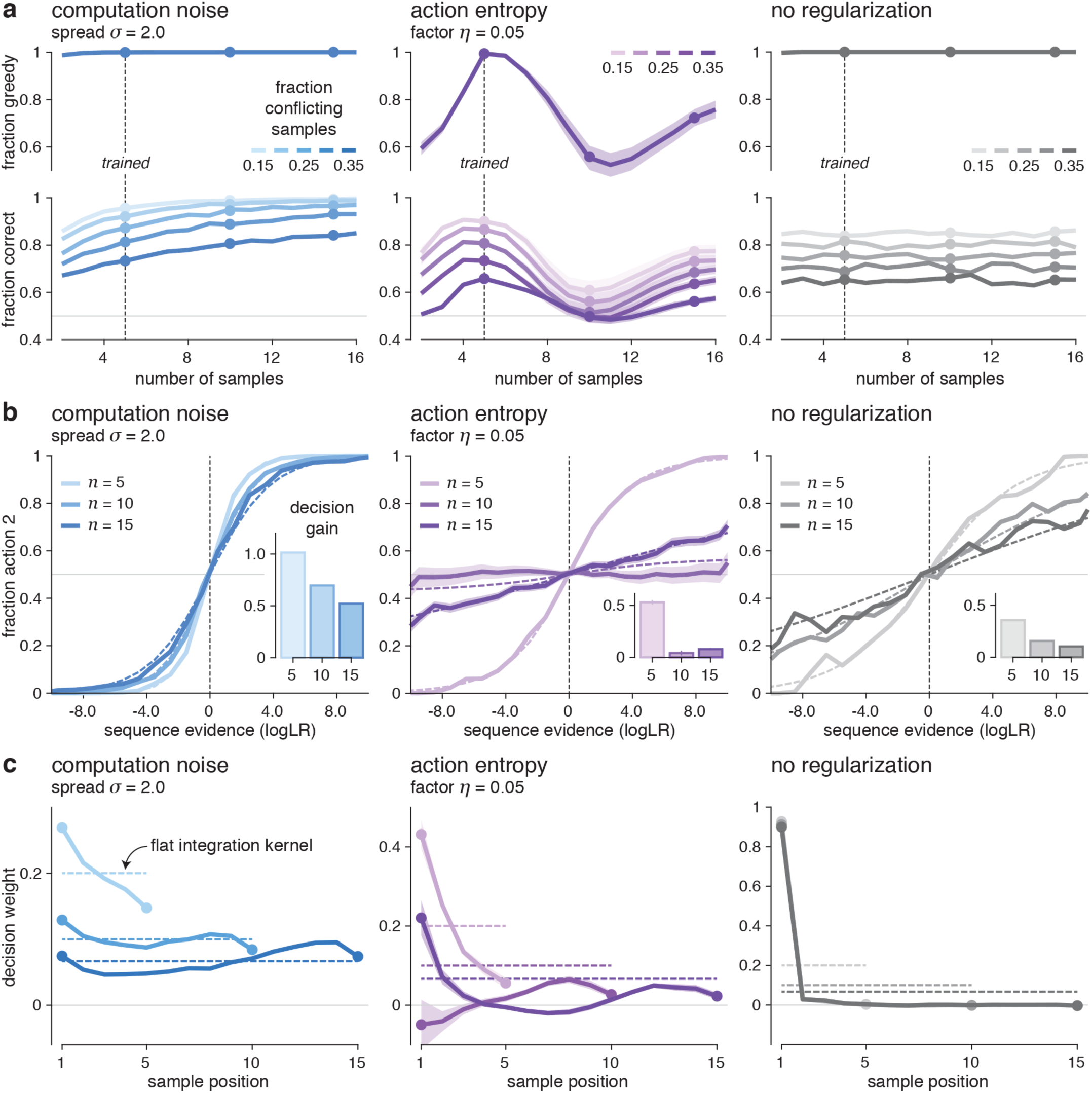
Signatures of Bayesian inference in the weather prediction task. **a**, Fraction of greedy decisions in stage 1 (top) and fraction of correct decisions in stage 2 (bottom) in response to sequences of 2 to 16 samples, for noisy RNNs (left; *σ* = 2), exact RNNs with action entropy regularization (middle; *η* = 0.05) and non-regularized RNNs (right). The accuracy of noisy RNNs grows monotonically with the number of samples in stage 2, an effect not shown by exact RNNs. **b**, Psychometric functions for noisy RNNs (left), exact RNNs with action entropy regularization (middle) and non-regularized RNNs (right). Noisy RNNs are highly sensitive to the Bayes-optimal evidence provided by each sequence. Inset: evidence sensitivity estimates for sequences of 5, 10 and 15 samples. **c**, Integration kernels for noisy RNNs (left), exact RNNs with action entropy regularization (middle) and non-regularized RNNs (right). Noisy RNNs integrate the provided evidence with moderate temporal biases. Dashed lines indicate flat (ideal) integration kernels.

We then estimated the sensitivity of noisy and exact RNNs to the Bayes-optimal evidence provided by each sequence (Fig. 3b). In contrast to exact RNNs, noisy RNNs were highly sensitive to this quantity across sequences of varying lengths. Furthermore, the estimation of the integration kernels used by noisy and exact RNNs through logistic regression (Fig. 3c; see Methods) showed that noisy RNNs were able to integrate the evidence provided by cues presented at successive samples with only moderate temporal biases. By contrast, exact RNNs trained with explicit action entropy regularization did not exhibit integration-compatible kernels, and non-regularized RNNs used only the first sample to decide – an extreme ‘primacy’ bias. Together, these findings indicate that training noisy RNNs in stage 1 endowed artificial agents with the ability to perform Bayesian inference, and thus to carry out stage 2 with high accuracy.

Inspired by recent observations in humans^18^, we performed additional quantitative analyses of the noisy RNNs to decompose their errors in stage 2 into two distinct sources: 1. errors triggered by computation noise, and 2. errors triggered by the action policy used by the RNN (Supplementary Fig. 1). These analyses revealed that the errors made by noisy RNNs, like humans, are mostly triggered by computation noise (Supplementary Fig. 1a). We also validated that, as in humans, the computation noise present in the network accumulates across samples of the same sequence – even when considering the non-flat integration kernels used by the network (Supplementary Fig. 1b). Finally, by comparing the actions taken by each noisy RNN to two repetitions of the same sequence of cues, we split the errors made by noisy RNNs in terms of a ‘bias-variance trade-off’ (Supplementary Fig. 1c). The bias terms reflects systematic deviations from Bayesian inference, whereas the variance term reflects random deviations due to computation noise. As in humans, the accuracy of noisy RNNs in stage 2 is bounded by the variance term – i.e., the limited effective precision of Bayesian inference performed by the artificial agents – and not by large systematic deviations (e.g., heuristics) from Bayesian inference.

### Computation noise reflects cue reliability in noisy agents

After studying the behavior of noisy and exact RNNs, we turned to the analysis of their activity patterns using dimensionality reduction techniques. First, we examined the input weights associated with each cue (Supplementary Fig. 2). Noisy and exact RNNs diverged very strongly in how they encoded the eight cues at the input level. Indeed, noisy RNNs projected all eight cues on a line, with cues associated with the same action projected in the same direction and cues associated with different actions projected in opposite directions (Supplementary Fig. 2a). In agreement with this line-based encoding of cues in noisy RNNs, the first principal component of input weights captured more than 90% of their overall variance across cues. By contrast, exact RNNs did not project cues on a line, with the first principal component of input weights capturing only 25% of their overall variance. Furthermore, the projection of the eight cues on the first principal component of input weights revealed an ordering of the cues as a function of their reward probabilities (likelihoods in Bayesian terms) in noisy RNNs (Supplementary Fig. 2b) – not in exact RNNs.

We then studied the statistics of activity patterns triggered by each cue in stage 1 (Fig. 4). In noisy RNNs, the projection of cue-wise activity patterns onto their mean direction showed a variability which decreases with their reliability (Fig. 4a). In other words, the depth of the ‘potential well’ associated with each cue increases with its reliability – making a more reliable cue more stable in terms of its activity pattern than a less reliable cue. Importantly, the standard deviation of projected activity patterns increased by 80% from the most reliable to the least reliable cues, whereas their mean only decreased by 8% (Fig. 4b). This observation indicates that it is the variability of activity patterns (triggered by computation noise) which encodes the reliability of individual cues in noisy RNNs, not their mean activation levels. This observation remained true for any presentation time of each cue (Supplementary Fig. 3a).

**Fig. 4.**
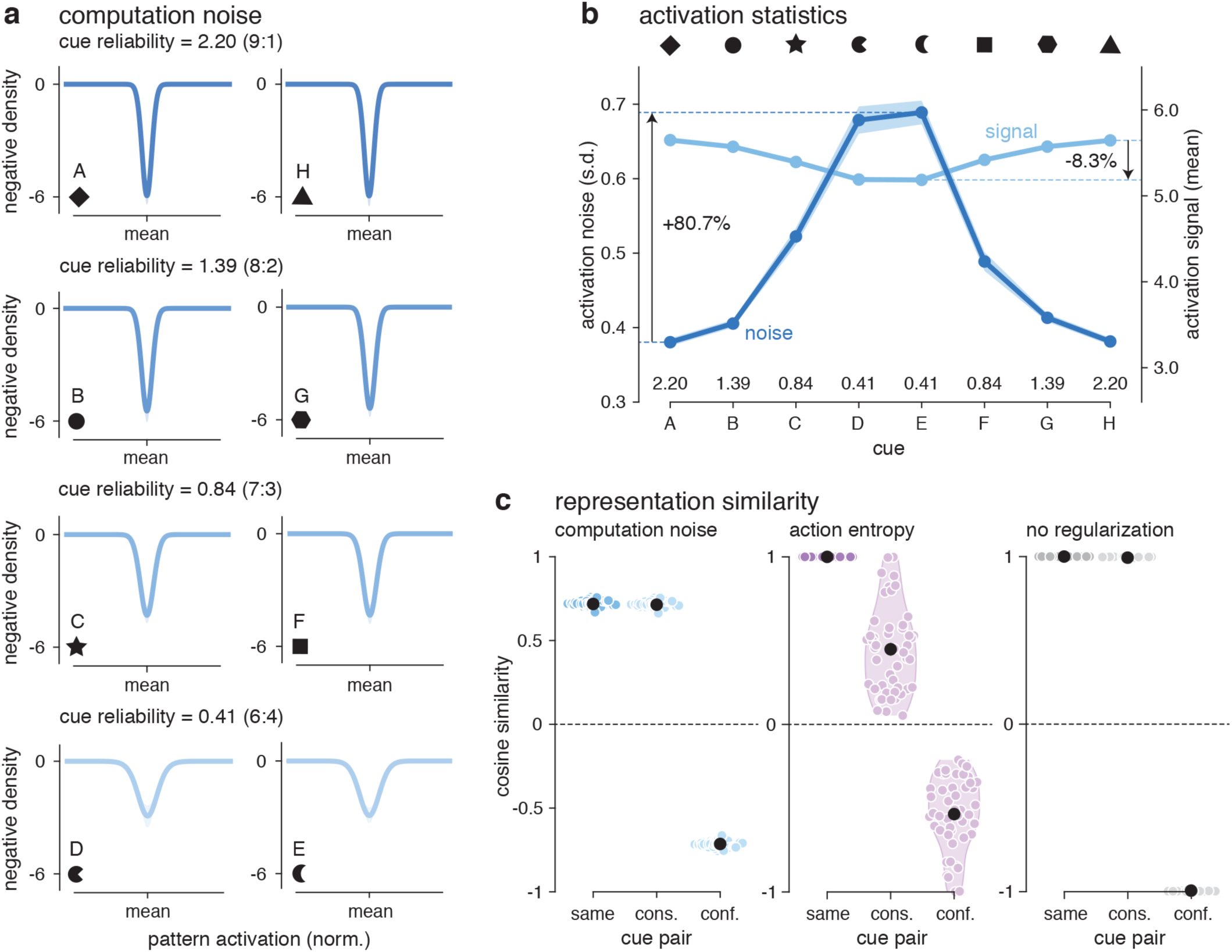
Encoding of cue reliability by computation noise. **a**, Effective potential wells associated with the 8 cues as a function of their reliability (top: most reliable cues; bottom: least reliable cues) in stage 1. The variability of cue-wise activity patterns projected onto their mean directions decreases with cue reliability. **b**, Activation statistics (light line: mean; dark line: standard deviation) associated with the 8 cues. The standard deviation of projected activity patterns increased by 80% from the most reliable to the least reliable cues, whereas their mean only decreased by 8%. **c**, Representation similarity analysis for noisy RNNs (left; *σ* = 2), exact RNNs with action entropy regularization (middle; *η* = 0.05) and non-regularized RNNs (right). Cosine similarity between activity patterns triggered by the same cue (left), by two ‘consistent’ cues associated with the same rewarded action (middle), and by two ‘conflicting’ cues associated with different rewarded actions (right).

Consistent with the line-based encoding of cues in noisy RNNs, the similarity between activity patterns triggered by different cues associated with the same action was as high as the similarity between activity patterns triggered by the same cue (Fig. 4c and Supplementary Fig. 3b). This was not the case for exact RNNs trained with action entropy regularization. Non-regularized RNNs showed in stage 1 only two precisely opposite activity patterns – one associated with each action. Together, these analyses indicate that noisy RNNs encode the reliability of individual cues in the trial-to-trial variability of their activity patterns. In other words, computation noise appears to have a functional role during testing, beyond regularization during training of the network weights.

### Computation noise knock-out disrupts Bayesian inference

If computation noise bears a functional role in the ability of noisy RNNs to perform Bayesian inference in stage 2, then knocking out computation noise after training of the network weights should impair the hallmarks of Bayesian inference identified in ‘wildtype’ RNNs (with computation noise during both training and testing). By contrast, if the main function of computation noise is to regularize the training of the network weights (as dropout^36^, for example), then knocking out computation noise during testing should either improve or not affect accuracy.

Noise ‘knocked-out’ RNNs performed equally well as noisy RNNs (i.e., optimally) when tested in stage 1 (Fig. 5a, top). By contrast, the accuracy of knocked-out RNNs in stage 2 (weather prediction) was substantially worse than wildtype RNNs (Fig. 5a, bottom). And importantly, the accuracy of knocked-out RNNs did not increase anymore with the number of presented samples. Even when only 5 samples were presented in stage 2 as during training (Fig. 5b), knocked-out RNNs performed no better than an agent relying on the first cue to decide (the extreme primacy bias exhibited by nonregularized RNNs). And indeed, the integration kernels of knocked-out RNNs (Fig. 5c) resembled the extreme primacy kernels estimated for non-regularized RNNs. This selective knock-out of computation noise during testing shows that, unlike dropout, computation noise is necessary even after training of the network weights for the network to perform Bayesian inference.

**Fig. 5.**
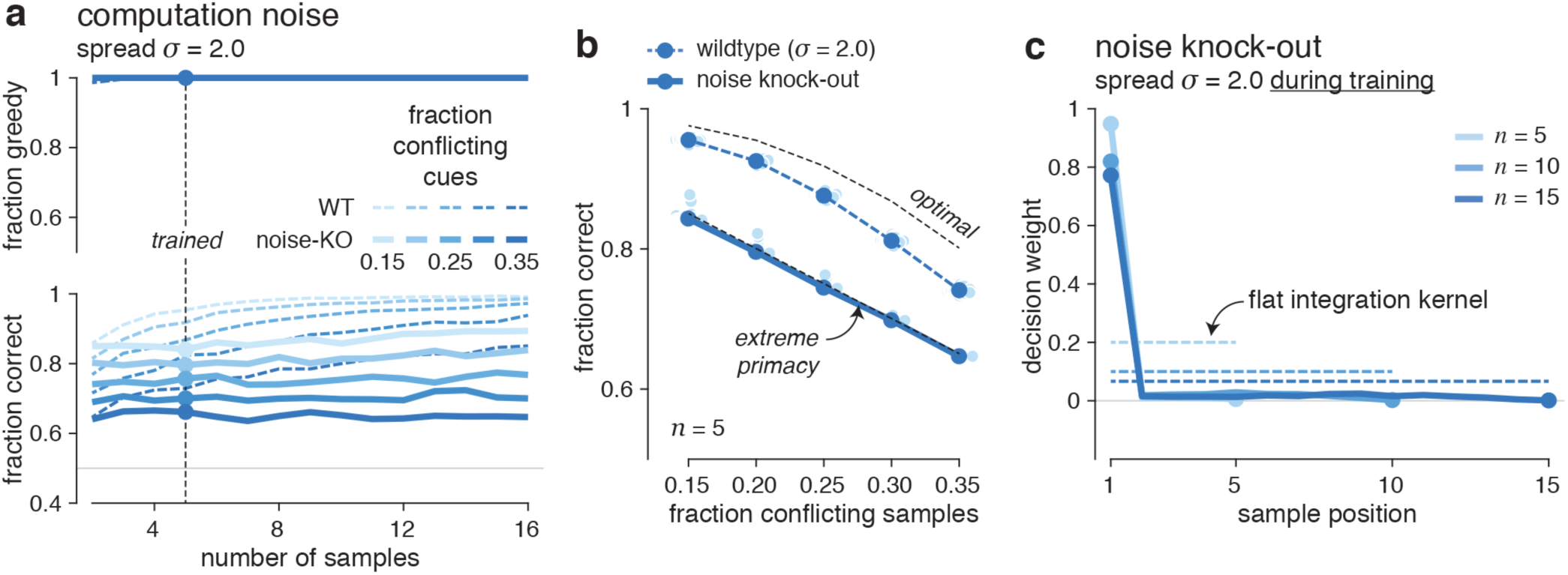
Computation noise knock-out in the weather prediction task. **a**, Fraction of greedy decisions in stage 1 (top) and fraction of correct decisions in stage 2 (bottom) in response to sequences of 2 to 16 samples, for ‘wildtype’ noisy RNNs (dashed lines; *σ* = 2) and ‘noise knocked-out’ RNNs (unbroken lines; *σ* = 2 during training and *σ* = 0 during testing). The accuracy of noise knocked-out RNNs does not grow with the number of samples in stage 2. **b**, Fraction of correct decisions in stage 2 (*n* = 5 samples) as a function of task difficulty for wildtype noisy RNNs (dashed line) and noise knocked-out RNNs (unbroken line). The black dashed lines indicate the accuracy of the Bayes-optimal decision-maker and the accuracy of a decision-maker using only the first sample (extreme primacy). **c**, Integration kernels for noise knocked-out RNNs for sequences of 5, 10 and 15 samples. Dashed lines indicate flat (ideal) integration kernels. Like exact RNNs with no regularization during training, noise knocked-out RNNs show an extreme primacy bias.

### Orthogonal encoding of likelihoods and posterior beliefs in noisy agents

The behavior of noisy RNNs and their line-based representation of individual cues at the input level are both consistent with Bayesian inference. But are activity patterns in stage 2 truly consistent with the latent variables computed during Bayesian inference? To address this question, we studied the two first principal components of activity patterns in noisy and exact RNNs during the presentation of 10^3^ sequences of cues in stage 2 (Fig. 6a; see Methods). The sequential Bayesian inference process required for the weather prediction task (stage 2) relies on the computation of two key variables: 1. the likelihood associated with the current cue, and 2. the posterior belief which integrates likelihoods across cues up to the current cue. These two variables have widely different time scales: the likelihood varies rapidly from one cue to the next, whereas the posterior belief integrates likelihoods over time and thus decays much less rapidly.

**Fig. 6.**
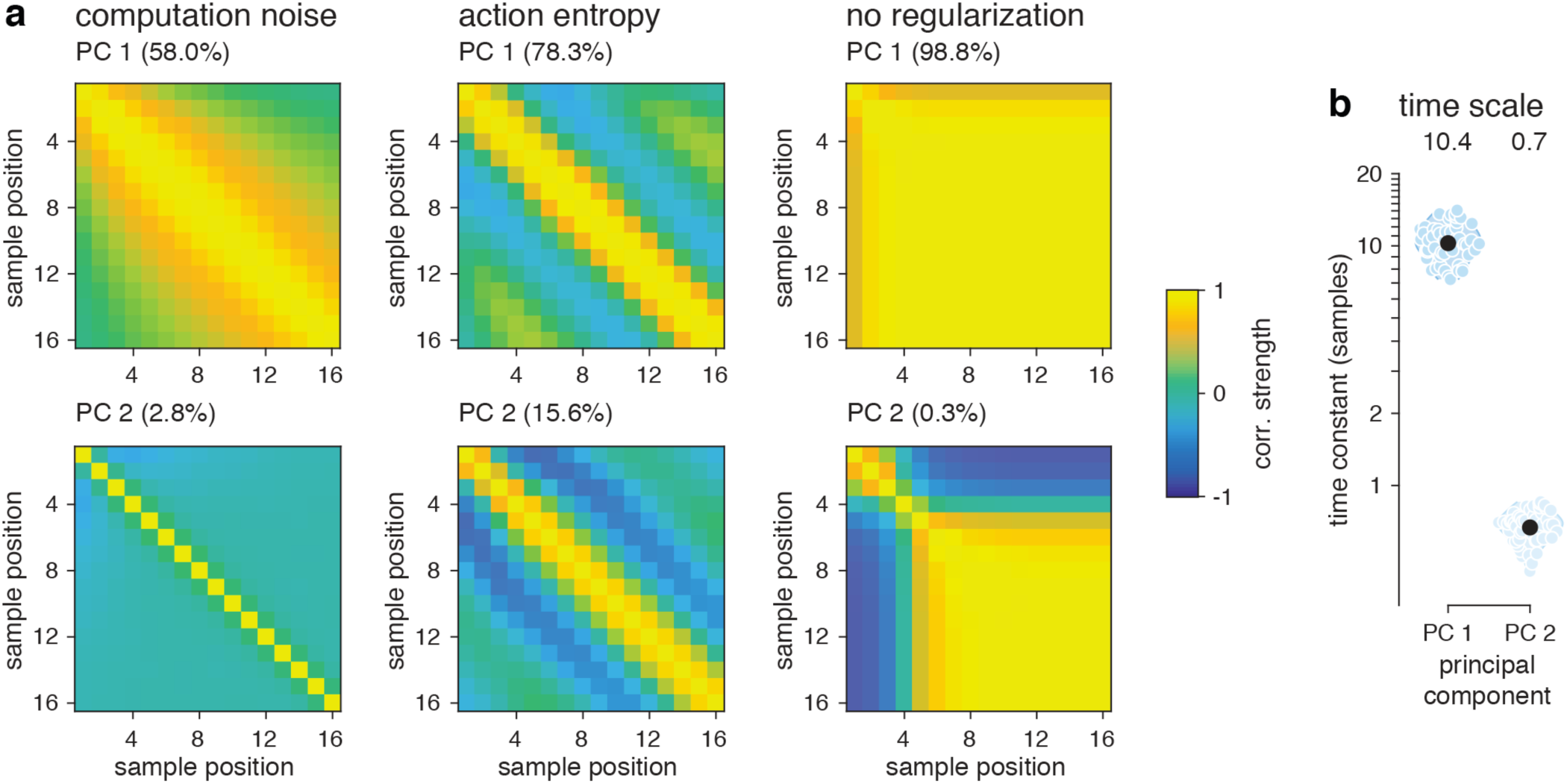
Principal component analysis of activity patterns in the weather prediction task. Each agent (computation noise: *σ* = 2; action entropy regularization: *η* = 0.05) is simulated on 1,000 trials of the weather prediction task (*n* = 16 samples; 25% of conflicting samples). **a**, Temporal autocorrelation matrices associated with the two first components (top: PC 1; bottom: PC 2) of noisy RNNs (left), exact RNNs with action entropy regularization (middle) and non-regularized RNNs (right). The two first PCs of activity patterns in noisy RNNs match the time scales of the two latent variables predicted by Bayesian inference (posterior belief and likelihood). **b**, Estimated time scales (exponential decay of autocorrelation coefficients) of the two first PCs of activity patterns in noisy RNNs. The time scale of PC 1 is of the order of 10 samples, whereas the time scale of PC 2 is shorter than 1 sample.

The two first principal components of activity patterns in noisy RNNs – orthogonal by construction – matched very well the time scales of the two latent variables predicted by Bayesian inference (Fig. 6a). The first principal component (PC 1, 58.0% of the overall variance) showed a slowly decaying autocorrelation across successive time samples (Fig. 6a, top), whereas the second principal component (PC 2, 2.8% of the overall variance) showed a rapidly decaying autocorrelation across time (Fig. 6a, bottom). Exact RNNs did not exhibit a similar segregation of time scales between their first PCs. In noisy RNNs, the time scale (autocorrelation decay) of PC 1 was of the order of 10 samples, whereas it was shorter than 1 sample for PC 2 (Fig. 6b).

PC 1 did not only show a time scale compatible with Bayesian inference: it followed the time course of the ideal Bayesian posterior belief in individual sequences, even those showing non-monotonic profiles (Fig. 7a). The correlation of PC 1 and PC 2 activities with likelihoods and posterior beliefs confirmed that PC 1 encodes the posterior belief (Fig. 7b, top), whereas PC 2 encodes the likelihood associated with the current cue (Fig. 7b, bottom). The temporal cross-correlation between PC 1 activity and the posterior belief further showed that noisy RNNs track the posterior belief with zero lag in PC 1 activity (Fig. 7c), a feature not shared with exact RNNs. Plotting the mean activity trajectories of noisy and exact RNNs in the two-dimensional space defined by their first PCs provided additional insight (Supplementary Fig. 4). The activity of noisy RNNs converged rapidly toward highly consistent endpoints across trained agents, endpoints whose distance from zero scaled with the number of presented samples. By contrast, the activity of exact RNNs trained with explicit action entropy regularization showed wild swings across the activity space over time, and the activity of non-regularized RNNs converged toward fixed endpoints independent of the number of presented samples – neither of these activity trajectories being compatible with Bayesian inference.

**Fig. 7.**
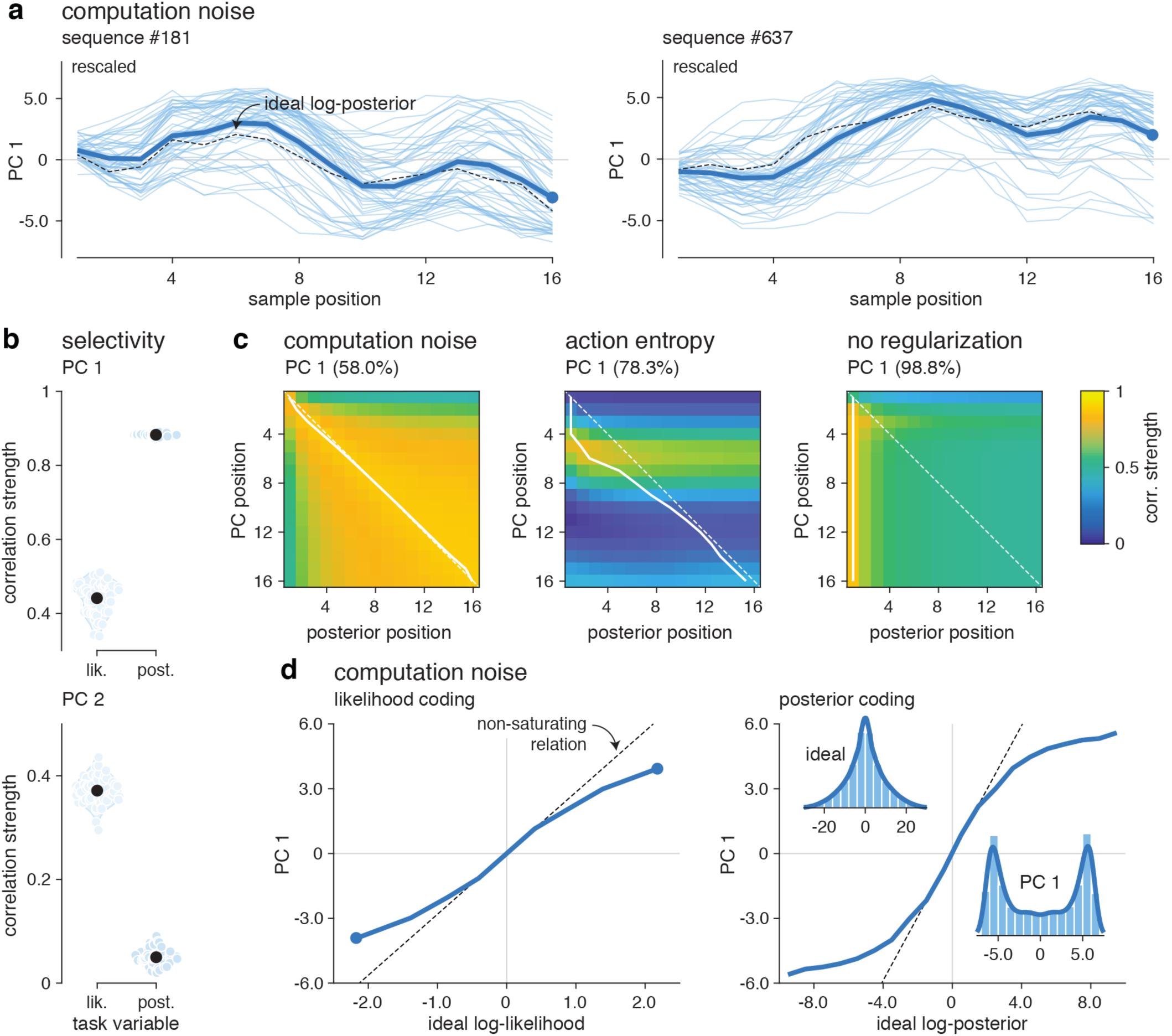
Orthogonal encoding of likelihoods and posterior beliefs in noisy RNNs. **a**, Time course of PC 1 for two representative sequences in stage 2 (*n* = 16 samples) presented to the 50 noisy RNNs with *σ* = 2. The black dashed line indicates the time course of the ideal log-posterior (rescaled for display purposes), whereas the blue unbroken line indicates the mean time course of PC 1 across the 50 noisy RNNs. Thin lines indicate the time course of PC 1 for each noisy RNN. **b**, Selectivity of PC 1 (top) and PC 2 (bottom) to log-likelihoods (left) and log-posterior beliefs (right). PC 1 activity correlates almost perfectly with the posterior belief, whereas PC 2 activity correlates only with the log-likelihood associated with the presented cue. **c**, Temporal cross-correlation matrices between the ideal log-posterior (x-axis) and PC 1 activity (y-axis) for noisy RNNs (left), exact RNNs with action entropy regularization (middle), and non-regularized RNNs (right). The thick white line indicates the correlation lag between PC 1 activity and the ideal log-posterior, showing that noisy RNNs encode the ideal log-posterior with near-zero lag (indicated by the white dashed line). **d**, Relations between PC 1 activity and the ideal log-likelihood (left), and with the ideal log-posterior (right). PC 1 activity correlates near-linearly with the ideal log-likelihood, whereas it shows a saturating relation with the ideal log-posterior. Inset: empirical probability distributions for the ideal log-posterior (top left) and PC 1 activity (bottom right) across trials and sample positions.

Finally, we plotted the precise relations between PC 1 activity in noisy RNNs and the variables computed during Bayesian inference (Fig. 7d). The ideal log-likelihood associated with the current cue was reflected near-linearly in PC 1 activity, whereas the ideal log-posterior showed a saturating relation with PC 1 activity. This encoding of the log-posterior in PC 1 activity suggests that noisy RNNs do not integrate information linearly, but rather in an attractor-like fashion. And together, these different effects confirm that the dominant activity patterns in noisy RNNs reflect the two latent variables computed during Bayesian inference. The fact that none of these effects are present in exact RNNs (trained with or without explicit action entropy regularization) indicates that it is the presence of computation noise which endows noisy RNNs with the ability to perform Bayesian inference. This cognitive ability was acquired by noisy RNNs without any training in the task which requires it to maximize accuracy.

### Alternative training parameters fail to promote Bayesian inference in exact agents

We examined whether tweaking aspects of the training of network weights in stage 1 could encourage exact RNNs – without computation noise – to perform Bayesian inference in stage 2. First, we considered parameters of the training condition (Supplementary Fig. 5a). Instead of using a fixed number of time samples, we presented each cue for a variable number of time samples before probing exact RNNs for a choice. This modification did not improve the extreme primacy bias exhibited by non-regularized RNNs (Supplementary Fig. 5b). We also considered omitting cue presentation at each time sample with probability *p* = 0.5. This second modification reduced the severity of the primacy bias, but non-regularized RNNs trained using this alternative scheme remained substantially worse than noisy RNNs in stage 2 (Supplementary Fig. 5c).

Because exact RNNs trained with explicit action entropy regularization showed clear signs of overfitting, we tested whether reducing the number of training steps could improve the accuracy of exact RNNs in stage 2. Plotting the accuracy of noisy and exact RNNs in stage 2 throughout training of the network weights revealed synchronous improvements in stage 1 and stage 2 for noisy RNNs (Supplementary Fig. 6a). In other words, training a noisy neural network to associate individual cues with rewarded actions configured the network to perform Bayesian inference in a fully unsupervised fashion. By contrast, the accuracy of exact RNNs trained with explicit action entropy regularization showed an early rise followed by a progressive decline in stage 2 characteristic of overfitting. This late decline was particularly pronounced for certain action entropy factors, possibly due to a runaway interaction between the entropy and policy terms of the objective function (Supplementary Fig. 6b). Therefore, we verified that stopping training of the network weights at the peak of the early rise did not lead to exact RNNs capable of performing Bayesian inference in stage 2. Indeed, exact RNNs trained using the action entropy factor which yields the highest accuracy in stage 2 did not show hallmarks of Bayesian inference (Supplementary Fig. 6c), and failed to approach the accuracy of noisy RNNs for long sequences of cues (Supplementary Fig. 6d).

### Zero-shot adaptation to volatile environments in noisy agents

Our findings show that computation noise in RNNs enables zero-shot learning of the weather prediction task from previously acquired stimulus-response associations. The behavior and activity patterns of noisy RNNs showed clear signatures of Bayesian inference, which made noisy RNNs surprisingly resilient to adverse conditions unseen during training: 1. sequences of conflicting cues associated with different rewarded actions, and 2. sequences of arbitrarily large numbers of cues, which noisy RNNs took advantage of to improve their accuracy. To probe the cognitive resilience of noisy RNNs further, we studied how artificial agents adapted their behavior to a mid-sequence change in the rewarded action – and thus in the distribution of cues being presented (Fig. 8a). Such adaptation to ‘volatile’ environments is typically viewed as a sign of adaptive cognition and hierarchical inference. As before, we tested noisy and exact RNNs trained in stage 1 – i.e., in perfectly stable environments.

**Fig. 8.**
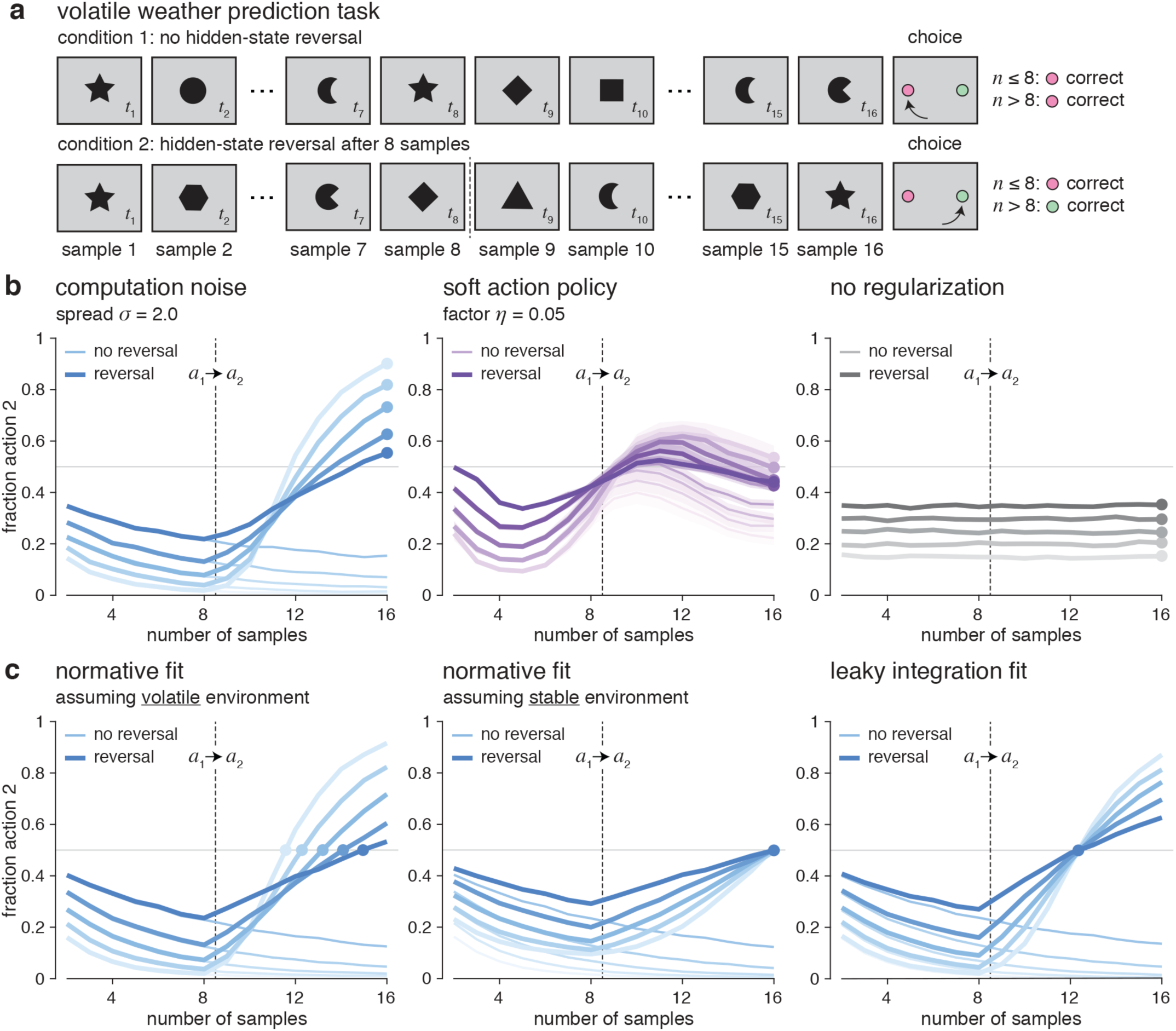
Volatile weather prediction task. **a**, Description of the experimental paradigm. In condition 1 (no reversal, top row), all samples are drawn from the same generative distribution associated with the same rewarded action. In condition 2 (reversal, bottom row), the first and last 8 samples are drawn from generative distributions associated with different rewarded actions. **b**, Fraction of trials where action 2 (the rewarded action for sequences longer than 8 samples) is chosen by noisy RNNs (left; *σ* = 2), exact RNNs with action entropy regularization (middle; *η* = 0.05) and non-regularized RNNs (right) in condition 1 (thin lines) and condition 2 (thick lines). In condition 2, noisy RNNs reverse their preferred action (from action 1 to action 2) in less than 8 trials on average, even for difficult sequences with a sizable fraction of conflicting samples (darker lines). **c**, Fits of normative integration (left: assuming non-zero volatility; middle: assuming zero volatility) and leaky integration (right) to the behavior of noisy RNNs. The adaptation of noisy RNNs to mid-sequence reversals is best explained by a Bayesian inference process assuming non-zero volatility.

In contrast to exact RNNs, noisy RNNs showed a remarkable ability to adapt their behavior to mid-sequence changes in the rewarded action (Fig. 8b). They reversed their action in less than 8 trials on average, even for difficult sequences with a sizable fraction of conflicting samples. Exact RNNs trained with action entropy regularization showed only a minimal adaptation of their behavior (riding above the large endogenous swings identified earlier), whereas non-regularized RNNs did not adapt their behavior at all to mid-sequence changes. We characterized the adaptation of noisy RNNs to mid-sequence changes by comparing two competing accounts: 1. a leaky integration process, and 2. a Bayesian inference process which assumes a non-zero volatility^37^ (i.e., the probability of change in the state of the environment). Fitting these two models to the behavior of noisy RNNs revealed that their adaptation to mid-sequence changes is best explained by a Bayesian inference process with non-zero volatility (Fig. 8c).

Plotting the mean activity trajectories of noisy and exact RNNs in trials featuring mid-sequence changes showed that noisy RNNs rapidly reconfigured their activity toward the endpoint defined by the rewarded action after the change (Fig. 9a). This was not the case for exact RNNs trained with action entropy regularization, whose endogenous dynamics were only minimally affected by the mid-sequence change, or for non-regularized RNNs whose activity did not move from the endpoint reached prior to the mid-sequence change. The time course of PC 1 activity following mid-sequence changes recapitulated the adaptability of noisy RNNs in volatile environments (Fig. 9b).

**Fig. 9.**
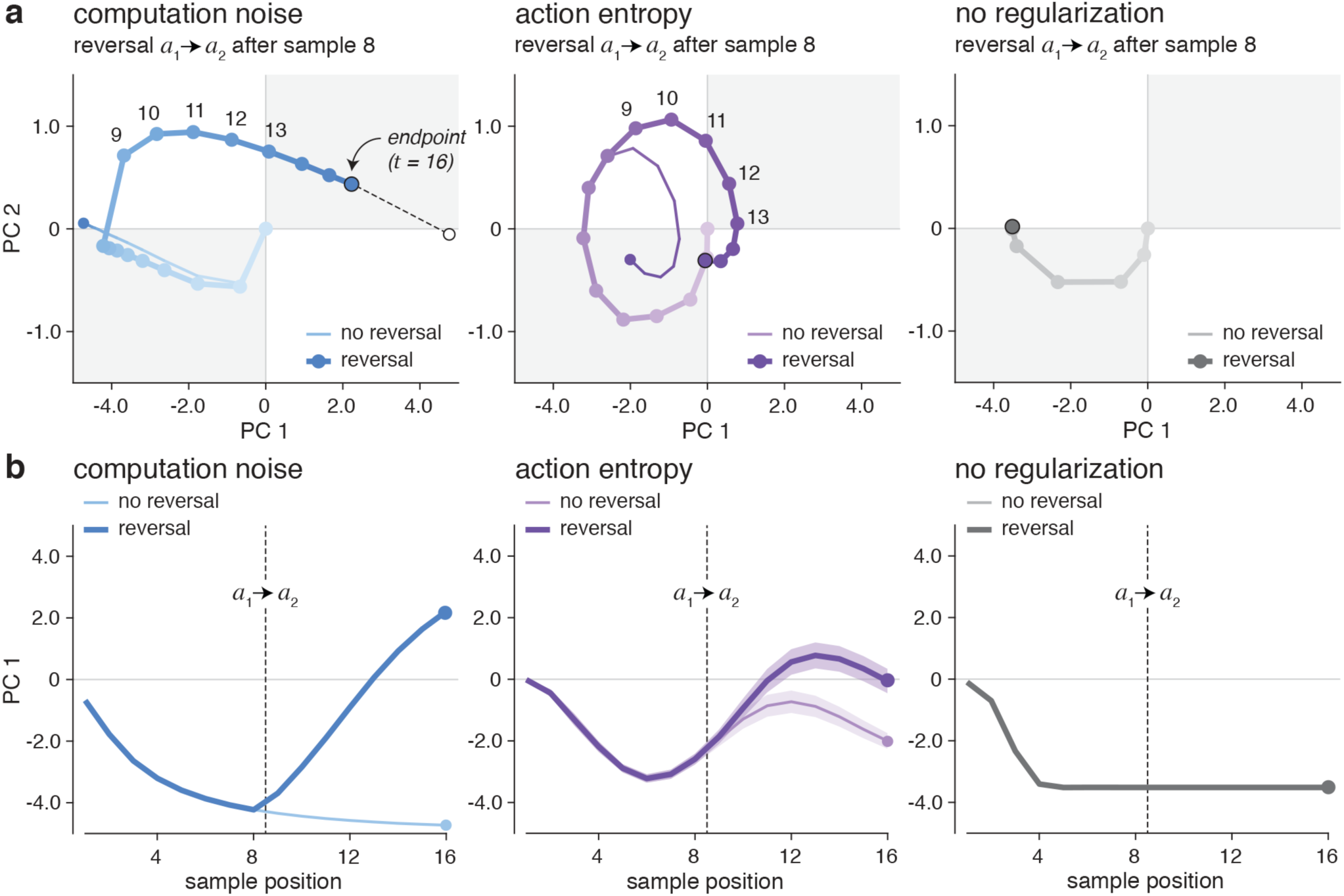
Trajectories of activity patterns in the volatile weather prediction task. **a**, Mean trajectories of activity patterns in the two-dimensional space defined by their first PCs for noisy RNNs (left; *σ* = 2), exact RNNs with action entropy regularization (middle; *η* = 0.05) and non-regularized RNNs (right). Thin lines indicate trajectories in condition 1 (no reversal), whereas thick lines indicate trajectories in condition 2 (reversal). Noisy RNNs rapidly reconfigure their activity toward the new endpoint defined by the rewarded action after the reversal. Exact RNNs trained with action entropy regularization show strong endogenous dynamics only minimally affected by the mid-sequence reversal. The activity of non-regularized RNNs does not move from the endpoint reached prior to the mid-sequence reversal. **b**, Time courses of PC 1 activity for noisy RNNs (left), exact RNNs with action entropy regularization (middle) and non-regularized RNNs (right). The first PC of noisy RNNs is compatible with the log-posterior of a Bayesian inference process assuming non-zero volatility.

Although noisy RNNs adapted their behavior to mid-sequence changes in the rewarded action, they did so relatively slowly – corresponding to a best-fitting volatility of 0.003, and thus an assumed change in the rewarded action every 300 samples on average. To determine whether noisy RNNs could adapt their behavior more rapidly in more volatile environments without any re-training of the network weights, we varied the input gain to the network (i.e., a simple multiplicative scaling of the input weights), and measured its impact on the adaptation of noisy RNNs to mid-sequence changes. Noisy RNNs adapted their behavior faster with increasing input gain, at the expense of worse asymptotic accuracy in sequences without mid-sequence changes (Supplementary Fig. 7a). These opposing effects of input gain on the behavior of noisy RNNs were well accounted for by the Bayesian inference process with non-zero volatility (Supplementary Fig. 7b). An increase in input gain was captured by a whopping 30-fold increase in the volatility assumed by the model – reaching 0.092 for an input gain of 4, which corresponds to an assumed change in the rewarded action every 10 samples on average (Supplementary Fig. 7c). Therefore, noisy RNNs trained to learn the probabilistic associations between isolated cues and rewarded actions are able to perform Bayesian inference near-optimally even in highly volatile environments.

### Lossless recovery from misleading outcomes in noisy meta-learning agents

Until now, the artificial agents we studied were trained to learn probabilistic stimulus-response associations through reinforcement learning, but did not update their representations of these associations when performing the weather prediction task – only combined them through passive sampling without interacting with the outcomes of their decisions. To understand whether computation noise promotes similar cognitive abilities in ‘meta-learning’ agents that sample actively their environment to maximize rewards even after training of the network weights^32,38^, we considered a second class of tasks based on the sequential sampling of ‘two-armed bandits’ (Fig. 10a). On each trial of a game, the agent is presented with a slot machine with two levers to choose from. The agent receives a reward as a function of the reward probability associated with the chosen lever in the current game, which in the most canonical instance of the task remains fixed over the course of the game. The most rewarded lever varies randomly across games, such that the agent needs to learn which lever is most rewarded in every single game. The difficulty of the task can be manipulated by varying the difference between the reward probabilities associated with the two levers.

**Fig. 10.**
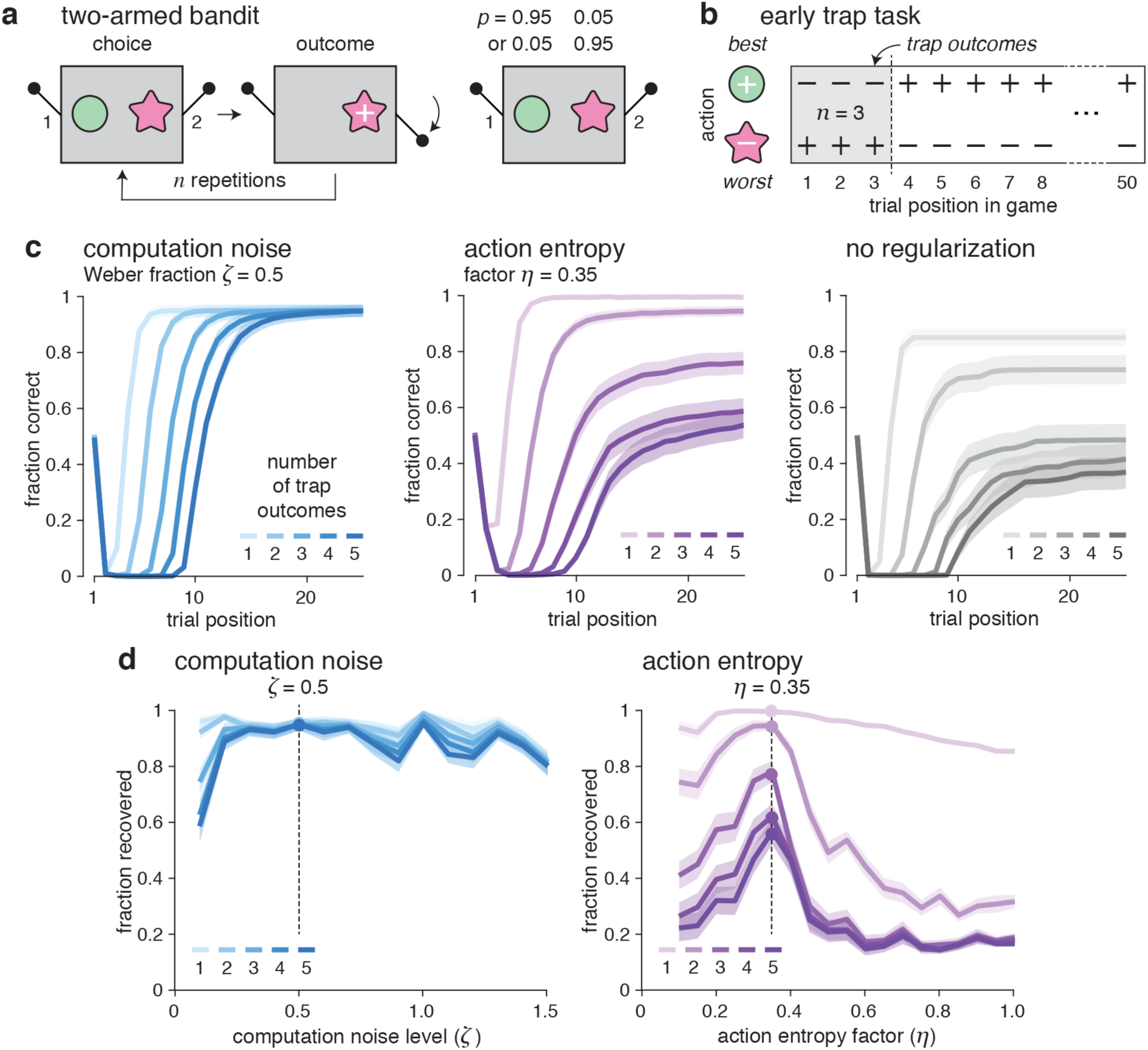
Meta-learning in the two-armed bandit task. **a**, Description of the experimental paradigm. On each trial of a game, the agent is presented with a slot machine with two levers to choose from. The agent receives a reward as a function of the reward probability associated with the chosen lever in the current game. The most rewarded lever varies randomly across games such that the agent needs to learn which lever is most rewarded in every single game. Artificial agents are trained using fixed reward probabilities of 0.95 and 0.05 for the two levers. **b**, Description of the ‘early trap’ task, where a number of misleading outcomes (up to 5) is presented at the beginning of each game. Performance in this task is measured as the ability of the agent to ‘recover’ from these early adverse events. **c**, Fraction of correct decisions (defined as trials where the agent chooses the most rewarded lever overall) in response to increasing numbers of early traps, for noisy RNNs (left; *ζ* = 0.5), exact RNNs with action entropy regularization (middle; *η* = 0.35) and non-regularized RNNs (right). Noisy RNNs recover almost systematically and independently from the number of early traps, whereas exact RNNs show clear signs of ‘jumping to conclusions’. **d**, Fraction of games where artificial agents have recovered from early traps, for noisy RNNs (left) and exact RNNs with action entropy regularization (right). Noisy RNNs recover substantially better from *n* > 1 early traps than exact RNNs across the whole range of tested computation noise levels and action entropy factors.

As before, we trained RNNs featuring either noisy or exact computations to perform the twoarmed bandit task with fixed reward probabilities of 0.95 and 0.05 for the best and worst levers (see Methods). We then tested the ability of these artificial agents to recover from misleading outcomes, or ‘traps’, at the beginning of each game (Fig. 10b). This ‘early trap’ condition constitutes an adverse condition for trained agents, especially when the number of successive early traps increases. As before, we varied the level of computation noise (fixed across training and testing) in the noisy RNNs, and the amount of explicit action entropy regularization in the exact RNNs (see Methods). Based on recent observations of such noise in humans^19,20^, computation noise during meta-learning was implemented by corrupting the updates of each unit in the network with i.i.d. normally distributed noise whose standard deviation scales with the quantity of update itself. This signal-dependent noise structure follows Weber’s law, a fundamental property of intensity sensation prevalent across perceptual domains and in the magnitude of associated neural responses. Computation noise is thus parameterized by its scaling coefficient *ζ* – also referred to as its ‘Weber fraction’.

Although noisy and exact meta-learning agents performed equally well in the condition in which they were trained, noisy RNNs recovered substantially better from early traps – even 5 successive traps at the beginning of the game – than their exact counterparts (Fig. 10c). While noisy RNNs recovered almost systematically and independently from the number of early traps, exact RNNs exhibited clear signs of ‘jumping to conclusions’: they did not fully recover from two or more traps at the beginning of the game, even when they were trained with explicit action entropy regularization. Noisy RNNs recovered efficiently from early traps for a wide range of computation noise levels (Fig. 10d). By contrast, action entropy regularization improved the recovery of exact RNNs from early traps only for a narrow range of scaling factors – for which the accuracy of regularized RNNs was still largely outperformed by noisy RNNs. This first experimental manipulation indicates that noisy meta-learning agents are resilient to several misleading outcomes at the beginning of a game – a situation never encountered during training.

### Zero-shot learning of the reversal learning task in noisy meta-learning agents

Because noisy meta-learning agents recovered efficiently from misleading outcomes at the beginning of the game, we asked whether they could also adapt their behavior after a reversal in the reward probabilities associated with the two levers in the middle of the game. This ‘reversal learning’ task^39^ is a canonical paradigm for studying adaptive learning and decision-making (Fig. 11a). Behavior in this task can be summarized by two curves: 1. the ‘learning curve’ in the first half of the game reflects how fast and how accurately the agent is able to identify the most rewarding lever, and 2. the ‘reversal curve’ in the second half of the game reflects how much the agent is able to adapt its beliefs and behavior to an unexpected change in reward probabilities.

**Fig. 11.**
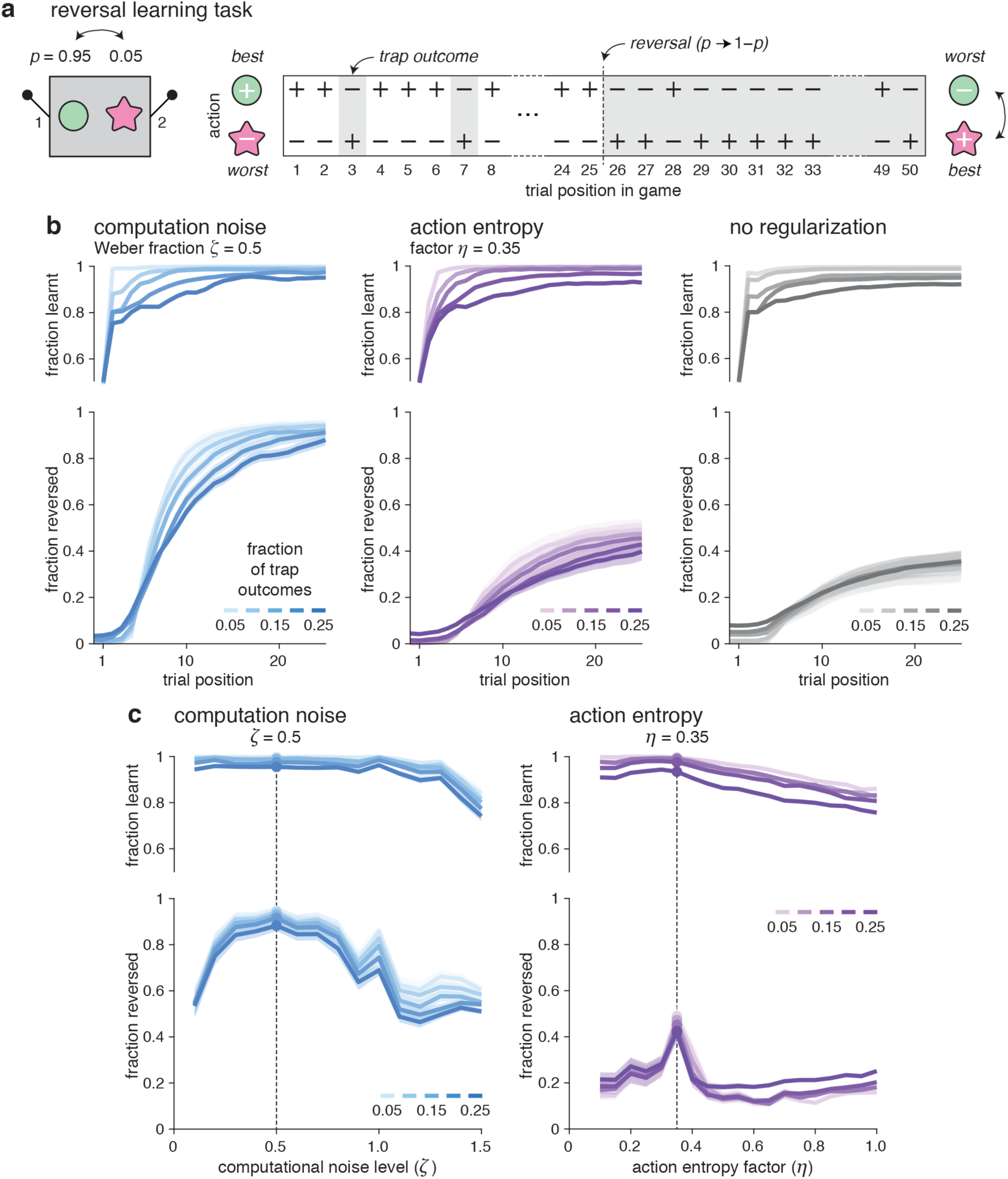
Reversal learning task. **a**, Description of the experimental paradigm. The reward probabilities associated with the two levers reverse in the middle of the game, such that the agent needs to switch away from a previously reinforced action. **b**, Learning curves (before reversal, top) and reversal curves (after reversal, bottom) during games with different fractions of trap outcomes, for noisy RNNs (left; *ζ* = 0.5), exact RNNs with action entropy regularization (middle; *η* = 0.35) and non-regularized RNNs (right). Learning curves do not differ between noisy and exact RNNs, whereas reversal curves show accurate adaptation to reversals only for noisy RNNs. **c**, Asymptotic learning rate (before reversal, top) and asymptotic reversal rate (after reversal, bottom) as a function of computation noise level in noisy RNNs (left), and of action entropy factor in exact RNNs (right). Noisy RNNs reverse efficiently their behavior for a wide range of computation noise levels, whereas action entropy regularization improves only moderately the behavior of exact RNNs.

Learning curves did not differ between noisy and exact RNNs (Fig. 11b, top). By contrast, reversal curves diverged sharply between the two types of agents: noisy RNNs reversed their behavior faster and substantially more often than exact RNNs – even those trained with action entropy regularization (Fig. 11b, bottom). As before, noisy RNNs reversed efficiently their behavior following the reversal in reward probabilities for a wide range of computation noise levels (Fig. 11c), whereas action entropy regularization moderately improved the reversal curve of exact RNNs only for a narrow range of scaling factors. In other words, noisy meta-learning agents are not only capable of recovering from early misleading outcomes, but also to switch away from an action which has been reinforced by positive outcomes for several trials.

Increasing the input gain to the network improved the ability of exact RNNs to reverse their behavior in the second half of the game, at the expense of worse initial learning in the first half (Supplementary Fig. 8a). But even exact RNNs driven by a performance-maximizing input gain failed to reverse their behavior nearly as efficiently as noisy RNNs (Supplementary Fig. 8b). In other words, the inflexibility of exact RNNs in the reversal learning task could not be compensated by increasing the input gain to the network.

### Computation noise supports adaptation to negative outcomes

To understand why noisy meta-learning agents are able to reverse their behavior much more efficiently than exact meta-learning agents trained in the same condition, we studied the statistics and dynamics of activity patterns in the two types of networks. First, we plotted the statistics of activity patterns associated with each action in the bandit task with fixed reward probabilities used to train the networks (Fig. 12a). The projection of action-wise activity patterns onto their mean direction after 50 trials of each game showed a larger trial-to-trial variability – and thus more shallow ‘potential wells’ – in noisy RNNs. This increased variability of activity patterns in noisy RNNs (Fig. 12b, left) was not only due to the intrinsic randomness associated with noisy activity updates in the network, but also by increased sensitivity of activity patterns to negative outcomes (Fig. 12b, right). Indeed, the presentation of only five negative outcomes decreased the dominant activity pattern in noisy RNNs by more than 60%, whereas it only did so by 20% or less in exact RNNs. The fact that the presentation of positive outcomes did not alter activity patterns in both types of networks betrays the presence of attractors (potential wells) associated with the two actions.

**Fig. 12.**
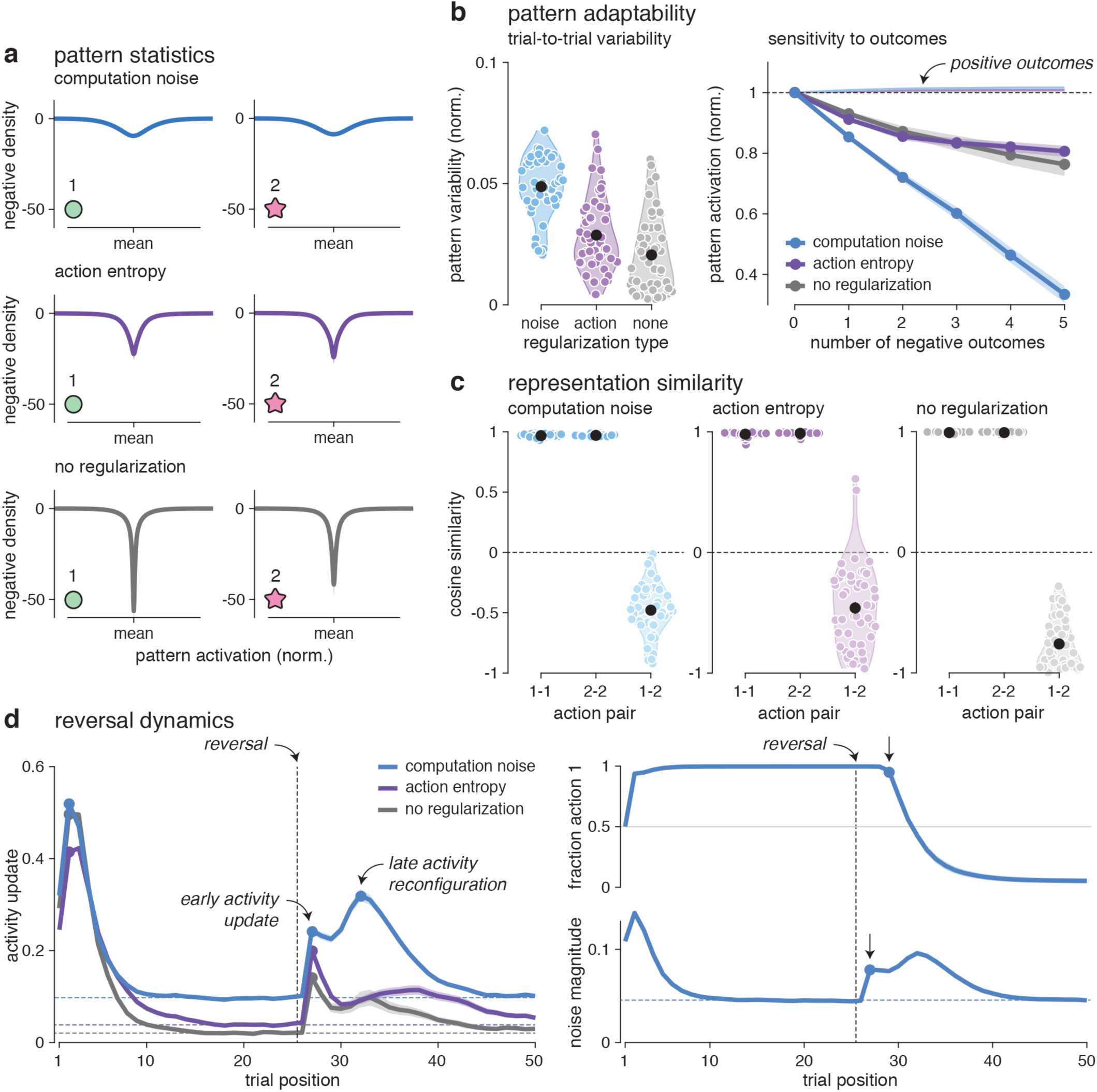
Destabilization of attractor states by computation noise. **a**, Effective potential wells associated with each action when reinforced by a fixed reward probability of 0.95, for noisy RNNs (top; *ζ* = 0.5), exact RNNs with action entropy regularization (middle; *η* = 0.35) and non-regularized RNNs (bottom). **b**, Pattern adaptability to negative outcomes. Left: pattern variability (standard deviation) associated with reinforced actions for noisy RNNs (left), exact RNNs with action entropy regularization (middle) and non-regularized RNNs (right). Right: destabilization of activation patterns by an increasing number of successive negative outcomes (up to 5). **c**, Representation similarity analysis for noisy RNNs (left), exact RNNs with action entropy regularization (middle) and non-regularized RNNs (right). Cosine similarity between activity patterns associated with the same reinforced action (left: 1-1; middle: 2-2) and by different reinforced actions (right: 1-2). **d**, Activity dynamics in the reversal learning task. Left: time course of activity update (unsigned difference between the recurrent activity at trial *t* and trial *t* − 1) for noisy (blue) and exact (violet and gray) RNNs. Right: time courses of choice behavior (top) and noise magnitude (bottom, defined as the unsigned difference between noisy and noise-free realizations of the update of recurrent activity at trial *t*).

The superiority of noisy RNNs over exact RNNs could not be explained by the similarity between the activity patterns associated with the two actions, but by the dynamics of activity in the two types of networks. While the two actions were associated with partially opposing activity patterns in both types of networks (Fig. 12c), noisy RNNs diverged strongly from exact RNNs in the update of their activity in the trials following a reversal (Fig. 12d, left). Both types of networks showed an early activity update in the first trial following the reversal – likely to be triggered by the first negative outcome associated with the previously reinforced action. But while this early activity update was transient in exact RNNs, noisy RNNs showed a second wave of activity update peaking 7 trials after the reversal – at the time when noisy RNNs switch toward the newly rewarded action for the first time. Aligning behavior with the magnitude of computation noise – which scales with trial-to-trial updates of activity in the network – showed that the increase in noise magnitude following the reversal precedes the change in behavior (Fig. 12d, right). Together, these results indicate that computation noise supports the enhanced behavioral flexibility of noisy meta-learning agents, by affording to destabilize existing activity patterns with few negative outcomes.

### Computation noise knock-out delays adaptation to adverse conditions

If computation noise bears a functional role for destabilizing attractor states in meta-learning agents, then knocking it out after training of the network weights should impair or at least delay the ability to adapt to adverse conditions – such as early traps (Fig. 10) or reversals in the middle of the game (Fig. 11). On the other hand, if computation noise serves primarily to regularize the training of the network weights, then knocking it out during testing should improve accuracy.

Like noisy RNNs, noise knocked-out RNNs could recover from early traps, but with delayed latency relative to wildtype RNNs (Fig. 13a). Across the whole range of tested computation noise levels, noise knocked-out RNNs never recovered better than wildtype RNNs – but substantially worse at higher computation noise levels (Fig. 13b). The effect of this selective knock-out of computation noise was highly similar for the reversal learning task: noise knocked-out RNNs adapted more slowly to reversals, but they learnt more rapidly the most rewarding lever in the initial learning phase (Fig. 13c). Similarly to the recovery from early traps, noise knocked-out RNNs never adapted better than wildtype RNNs to reversals across the whole range of tested computation noise levels (Fig. 13d). Together, these findings indicate that computation noise in meta-learning agents supports adaptation to adverse conditions even after training of the network weights.

**Fig. 13.**
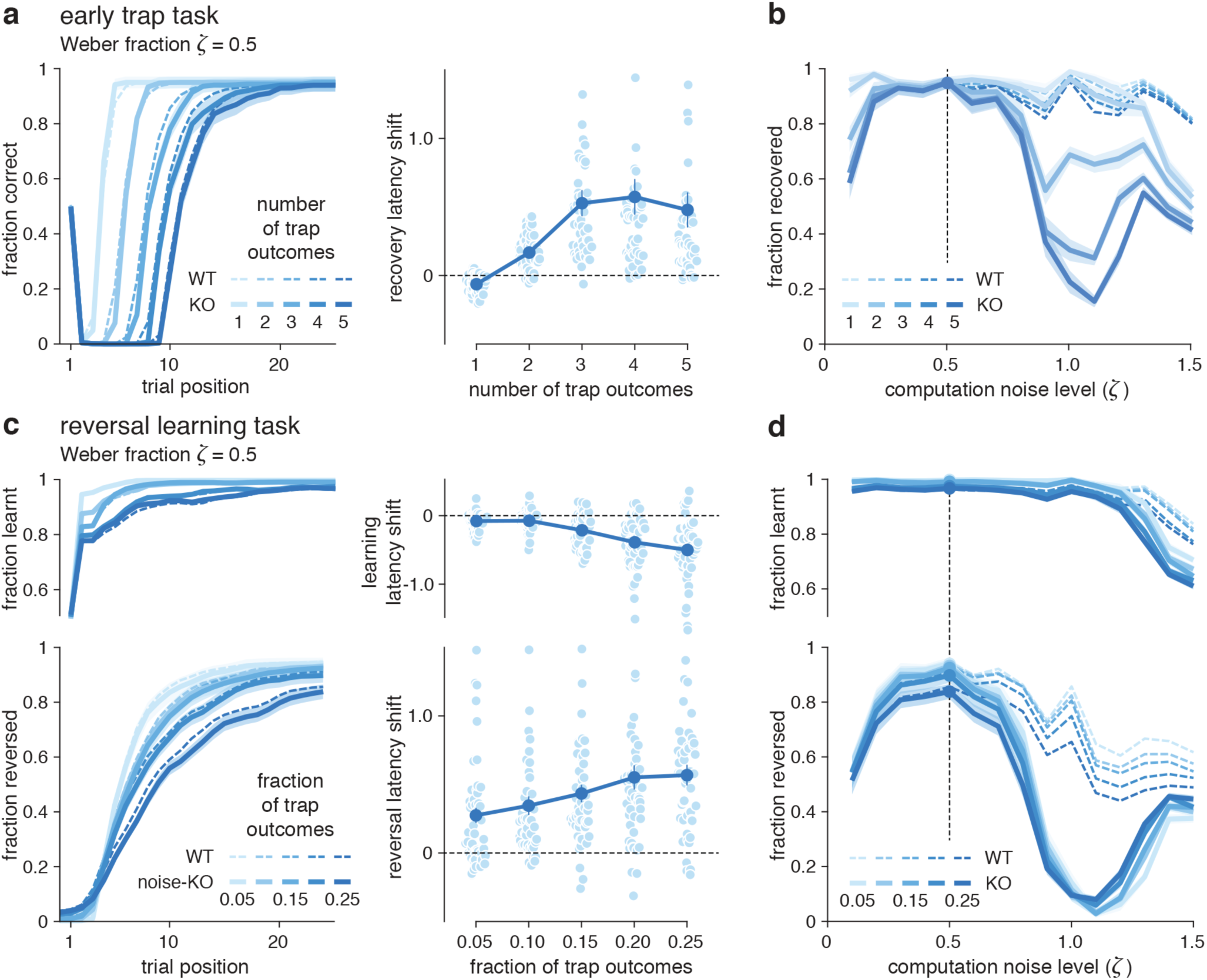
Computation noise knock-out in the bandit task. **a**, Early trap task. Left: fraction of correct decisions in response to increasing numbers of early traps, for ‘wildtype’ noisy RNNs (dashed lines; *ζ* = 0.5) and ‘noise knocked-out’ RNNs (unbroken lines; *ζ* = 0.5 during training and *ζ* = 0 during testing). Right: recovery latency shift triggered by computation noise knock-out (difference in recovery latency between noise knocked-out and wildtype RNNs). Noise knocked-out RNNs recover slightly more slowly than noisy RNNs. **b**, Fraction of games where artificial agents have recovered from early traps, for ‘wildtype’ noisy RNNs (dashed lines) and ‘noise knocked-out’ RNNs (unbroken lines). Computation noise knock-out impairs performance for large computation noise levels (*ζ* > 0.5). **c**, Reversal learning task. Left: learning curves (before reversal, top) and reversal curves (after reversal, bottom) for ‘wildtype’ noisy RNNs (dashed lines) and ‘noise knocked-out’ RNNs (unbroken lines). Right: learning latency shift (top) and reversal latency shift (bottom) triggered by computation noise knock-out. Noise knocked-out RNNs show slightly more rapid initial learning, but slower reversal learning than noisy RNNs. **d**, Asymptotic learning rate (before reversal, top) and asymptotic reversal rate (after reversal, bottom), for ‘wildtype’ noisy RNNs (dashed lines) and ‘noise knocked-out’ RNNs (unbroken lines). Computation noise impairs performance for large computation noise levels (*ζ* > 0.5).

### Orthogonal encoding of actions and outcomes in noisy meta-learning agents

What properties of activity patterns enable the more efficient adaptation to adverse conditions observed in noisy RNNs? To address this question, we studied the two first principal components of activity patterns in noisy and exact RNNs during the two-armed bandit task with fixed reward probabilities – i.e., the condition used to train the network weights (see Methods). For both types of networks, the two first principal components of activity patterns showed very different time scales (Fig. 14a). The first principal component (PC 1, more than 85% of the overall variance) showed high autocorrelation across trials (Fig. 14a, top), whereas the second principal component (PC 2, less than 5% of the overall variance) showed a rapid decay in autocorrelation across two successive trials (Fig. 14a, bottom). Nevertheless, autocorrelation matrices exhibited subtle differences between the two types of networks: while PC 2 activity did not correlate with itself across trials for noisy RNNs, residual autocorrelation patterns were apparent for exact RNNs.

**Fig. 14.**
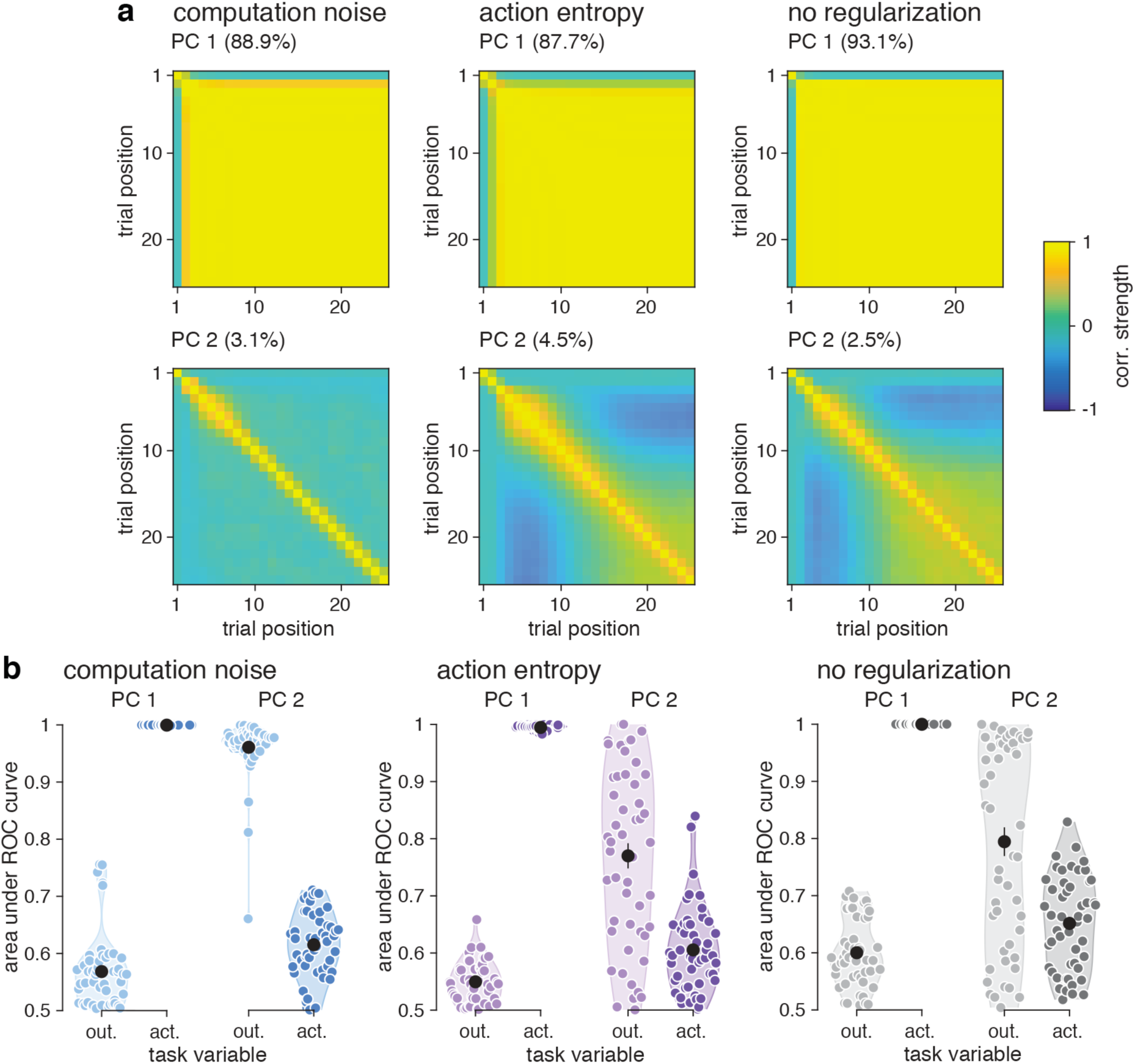
Principal component analysis of activity patterns in the bandit task. Each agent (computation noise: *ζ* = 0.5; action entropy regularization: *η* = 0.35) is simulated on 1,000 trials of the bandit task with fixed reward probabilities 0.95 and 0.05. **a**, Temporal autocorrelation matrices associated with the two first components (top: PC 1; bottom: PC 2) of noisy RNNs (left), exact RNNs with action entropy regularization (middle) and non-regularized RNNs (right). PC 1 shows a much stronger autocorrelation profile than PC 2, across all types of networks. PC 2 shows no autocorrelation across trials for noisy RNNs, not for exact RNNs. **b**, Selectivity of PC 1 and PC 2 to the current action (output) and the previous outcome (input), for noisy RNNs (left), exact RNNs with action entropy regularization (middle) and non-regularized RNNs (right). The double dissociation between the selectivity of the two PCs to task variables (PC 1: current action; PC 2: previous outcome) is particularly clear for noisy RNNs.

The correlation of PC 1 and PC 2 activities with the outcome obtained at the end of the previous trial and the action chosen in the current trial (Fig. 14b) revealed that PC 1 reflects the current action (i.e., the output of the network), whereas PC 2 reflects the previous outcome (i.e., the input to the network). This double dissociation between the selectivity of the two PCs to task variables was particularly clear for noisy RNNs, and less obvious for exact RNNs. Plotting the mean activity trajectories of noisy and exact RNNs in the two-dimensional space defined by their first PCs highlighted large differences during reversal learning (Fig. 15a). The two PCs were signed such that the most rewarding action before the reversal is associated with a positive PC 1 activity, and a positive outcome obtained at the end of the previous trial is associated with a positive PC 2 activity.

**Fig. 15.**
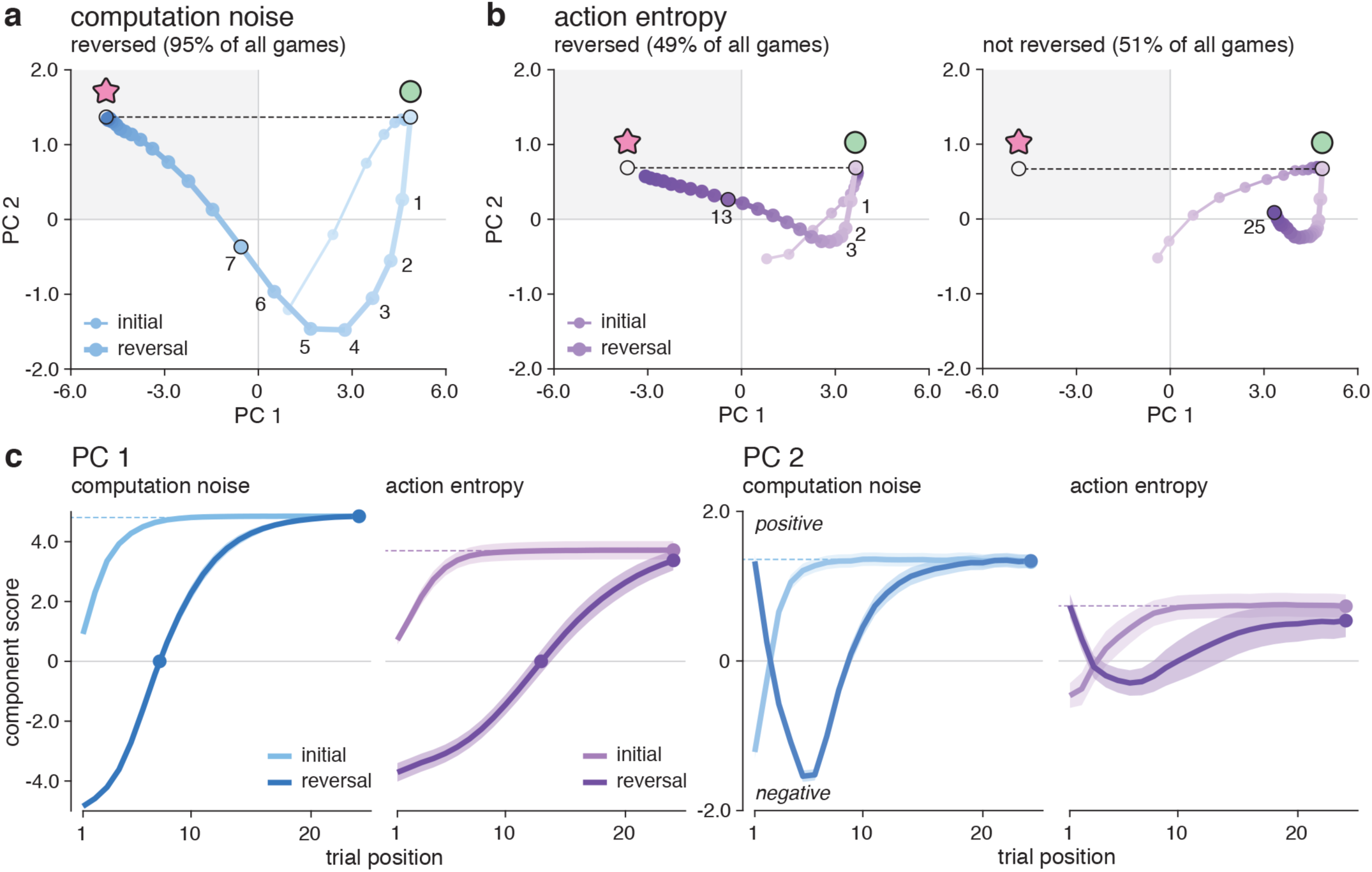
Trajectories of activity patterns in the reversal learning task. **a**, Mean trajectories of activity patterns in the two-dimensional space defined by their first PCs for noisy RNNs (*ζ* = 0.5) in reversal learning games where the network reversed its behavior (95% of all games). The thin line indicates the trajectory in the initial learning phase (before reversal), whereas the thick line indicates the trajectory in the reversal learning phase (after reversal). After a reversal, noisy RNNs show a rapid negative decline of PC 2 (outcome-selective) activity, followed by a rapid shift of PC 1 (action-selective) activity toward the newly rewarded action. **b**, Mean trajectories of activity patterns for exact RNNs with action entropy regularization (*η* = 0.35) in reversal learning games where the network reversed its behavior (left; 49% of all games) or missed the reversal (right; 51% of all games). Missed reversals are preceded by larger PC 1 activity in exact RNNs. **c**, Time courses of PC 1 activity (left) and PC 2 activity (right) for noisy RNNs (blue lines) and exact RNNs (violet lines) with action entropy regularization, in the initial learning phase (lighter colors) and the reversal learning phase (darker colors). Activity dynamics are nearly identical across the two phases for noisy RNNs, but slower (PC 1) and dampened (PC 2) during reversal learning for exact RNNs.

After a reversal, noisy RNNs showed a rapid negative decline of PC 2 (outcome-selective) activity, likely due to the negative outcomes obtained from choosing the previously reinforced action, followed by a rapid shift of PC 1 (action-selective) activity toward the newly rewarded action (Fig. 15a). The activity trajectory of exact RNNs following a reversal distinguished very clearly the games in which the network reversed its behavior (Fig. 15b, left) from those in which the network failed to reverse its behavior (Fig. 15a, right). In particular, the value of PC 1 (action-selective) activity before a reversal predicted whether exact RNNs would subsequently adapt their behavior. Indeed, missed reversals were preceded by larger PC 1 activity – consistent with overly strong attractor states in these networks. Activity updates of the two PCs confirmed this interpretation, by showing that missed reversals in exact RNNs were associated with weaker updates of PC 2 (outcome-selective) activity in the first trial following the reversal, and the absence of a second update of PC 1 (action-selective) activity, which otherwise peaks 10 trials later when the exact RNNs manage to reverse its behavior (Supplementary Fig. 9).

The side-by-side comparison between the time courses of PC 1 and PC 2 activity during initial learning and reversal learning summarized the differences between noisy and exact RNNs: their dynamics were nearly identical across the two learning phases for noisy RNNs, but not for exact RNNs (Fig. 15c). These analyses of activity patterns in noisy and exact meta-learning agents indicate that computation noise results in the representation of action values and action outcomes – two key variables in reinforcement learning – as orthogonal activity patterns. In adverse conditions, this noise drives efficient transitions between attractor states, allowing meta-learning agents to adapt to several adverse events – including sudden reversals of reward probabilities in the middle of a game – entirely unseen during training of the network weights.

### Zero-shot tracking of drifting reward probabilities in noisy meta-learning agents

To further explore the cognitive resilience of noisy meta-learning agents, we presented them with another variant of the bandit task which constitutes an even clearer case of adverse condition: the ‘restless’ bandit task^19,40^, in which the reward probabilities associated with the two levers drift randomly over the course of the game (Fig. 16a). Each game is made of several implicit reversals every time reward probabilities cross each other. We used the same noisy and exact RNNs as before – i.e., trained in a bandit task with fixed reward probabilities of 0.95 and 0.05 (Fig. 10a).

**Fig. 16.**
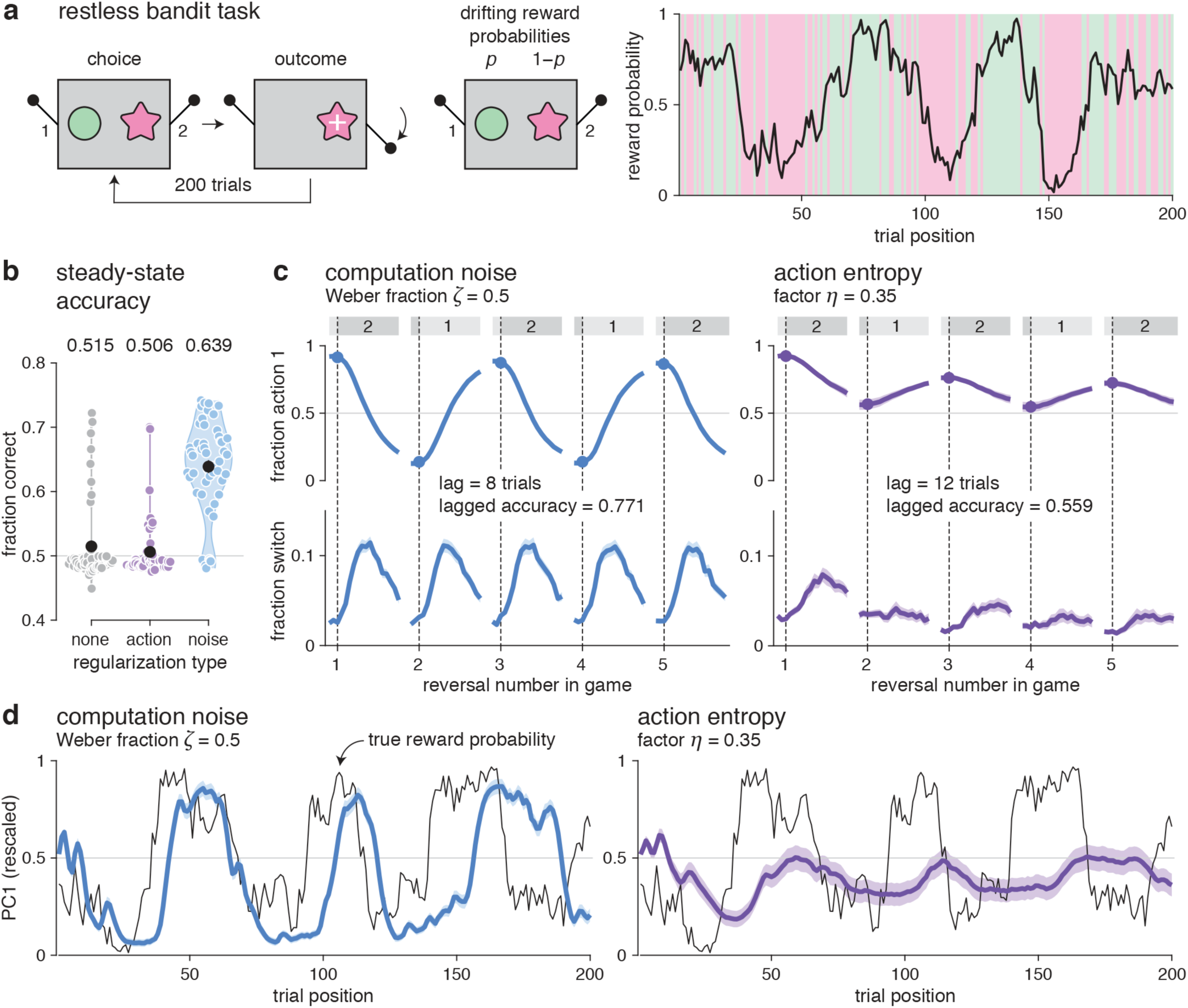
Restless bandit task. **a**, Description of the experimental paradigm. The reward probabilities associated with the two levers (*p* and 1 − *p*) drift randomly over the course of the game (200 trials). **b**, Steady-state accuracy (fraction of trials *t* where the network chooses the action with higher reward probability *p*_*t*_, after an initial lead-in period of 25 trials) for noisy RNNs (*ζ* = 0.5), exact RNNs with action entropy regularization (*η* = 0.35) and non-regularized RNNs. Only noisy RNNs are able to perform the restless bandit task above chance. **c**, Choice behavior following the first 5 implicit reversals of a game, for noisy RNNs (left) and exact RNNs with action entropy regularization (right). Top: fraction of trials where agents choose the most rewarding lever at the beginning of the game (labelled action 1). Bottom: fraction of trials where agents switch from one action at trial *t* − 1 to the other action at trial *t*. Noisy RNNs reverse their preferred action independently of the position of the reversal in the game, whereas exact RNNs remain biased toward the most rewarded action at the beginning of the game. The choice behavior of noisy RNNs lags by 8 trials behind reward probabilities, and by 12 trials for exact RNNs. **d**, Time course of PC 1 activity during a representative game, for noisy RNNs (left) and exact RNNs with action entropy regularization (right). PC 1 activity is rescaled to the range of simulated reward probabilities across 100 games for display purposes. The first PC of noisy RNNs tracks a filtered representation of reward probabilities, something not observed for exact RNNs.

Measuring the ‘steady-state’ accuracy of the two type of networks (i.e., the fraction of trials where the network chose the action with higher current reward probability, after an initial lead-in period of 25 trials) revealed that only noisy RNNs were able to perform the restless bandit task accurately (Fig. 16b). Even exact RNNs trained with explicit action entropy regularization performed no better than chance. The reversal learning curves associated with the first five implicit reversals of each game showed that noisy RNNs were able to reverse their behavior independently of the position of the reversal in the game, whereas exact RNNs became progressively insensitive to reversals over the course of the game (Fig. 16c). The behavior of noisy RNNs also lagged less behind reward probabilities (8 trials) than the behavior of exact RNNs (12 trials).

Examining PC 1 activity over time showed that noisy RNNs tracked a filtered representation of reward probabilities in their dominant (action-selective) activity pattern – something which was not found for exact RNNs trained and tested in the same conditions as noisy RNNs (Fig. 16d). In other words, computation noise does not only afford to adapt to sudden reversals in reward probabilities, but also to track continuous drifts in reward probabilities over extended periods of time – a cognitive ability highly reminiscent of Kalman filtering, the optimal Bayesian inference process for tracking reward probabilities in the restless bandit task.

### Zero-shot adaptation to volatile environments in noisy meta-learning agents

We have observed earlier that: 1. computation noise drives efficient transitions between attractor states following reversals, and 2. reversals are associated with a transient increase in the magnitude of computation noise in the network – owing to the signal-dependent structure of the noise. Based on these two observations, we hypothesized that computation noise could yield faster adaptation to reversals in conditions where reversals are more frequent – i.e., volatile environments. Such adaptation to volatility has typically been implemented either through explicit sophistication of cognitive models (e.g., a two-stage hierarchical inference process) or by training artificial neural networks in volatile environments. Here, by contrast, we used noisy and exact RNNs trained in a bandit task without any reversal – i.e., in perfectly stable environments.

To test this hypothesis, we compared noisy and exact RNNs in two conditions associated with different volatility^32,41^ – i.e., different reversal probability (Fig. 17a). As indicated by its name, the stable condition contained no reversal, whereas the volatile condition featured a non-zero volatility of 0.04 (in practice, a reversal every 25 trials). The probability of trap outcomes was fixed across conditions to 0.2. As in the reversal learning task (Fig. 11), noisy RNNs performed substantially better than exact RNNs in the volatile condition (Fig. 17b). But here, we further fitted to their behavior a normative inference model with two free parameters which correspond to the generative parameters of the task: 1. the inferred reversal probability (volatility), and 2. the inferred trap probability (see Methods).

**Fig. 17.**
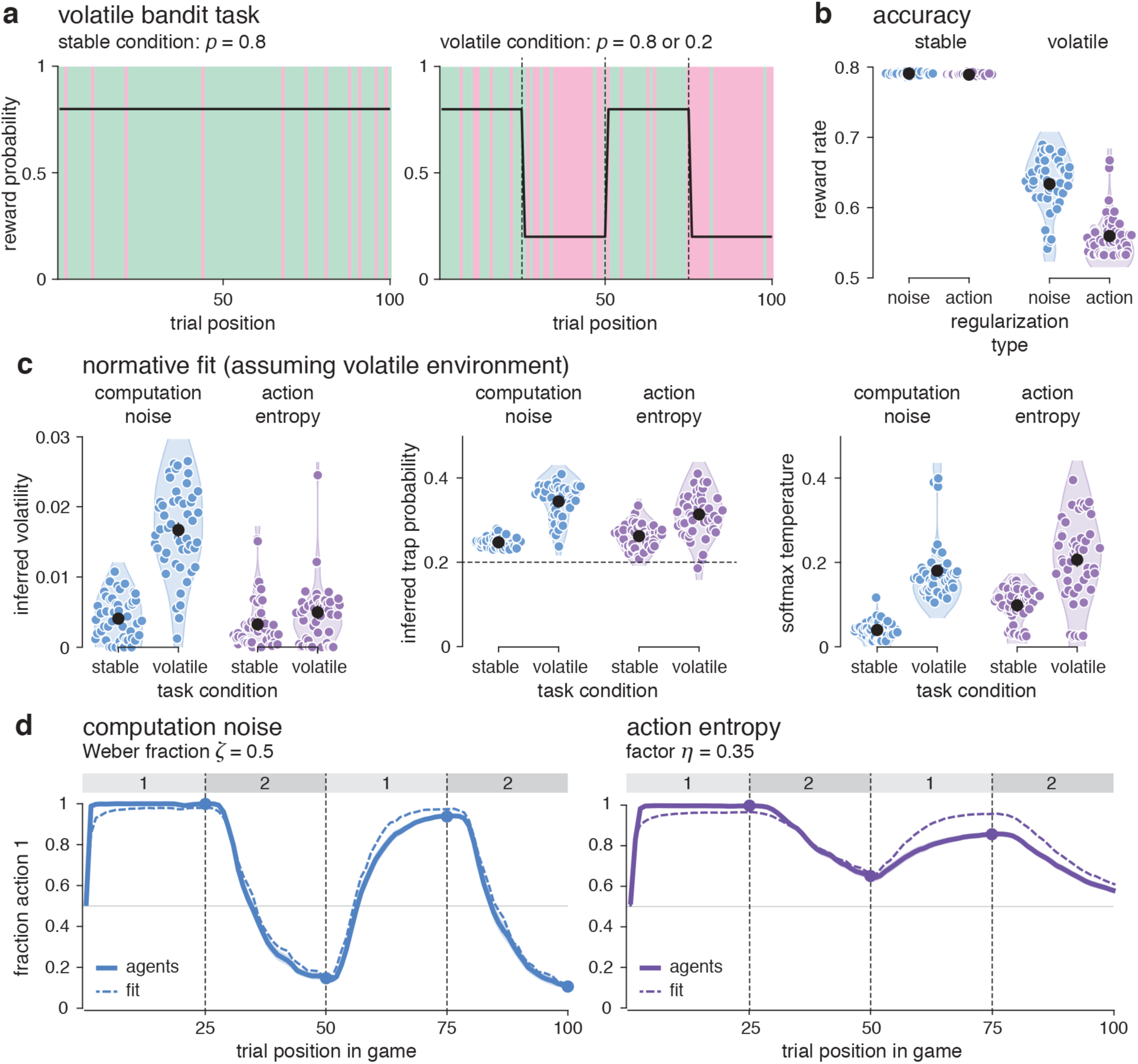
Volatile bandit task. **a**, Description of the experimental paradigm. Left: stable condition with fixed reward probabilities 0.8 and 0.2. Right: volatile condition with reversing reward probabilities every 25 trials. **b**, Reward rate achieved by noisy RNNs (*ζ* = 0.5) and exact RNNs with action entropy regularization (*η* = 0.35) in the stable (left) and volatile (right) conditions. Noisy and exact RNNs perform equally well (and optimally) in the stable condition, whereas noisy RNNs outperform exact RNNs in the volatile condition. **c**, Best-fitting estimates of inferred volatility (left), inferred trap probability (middle) and softmax temperature (right), for noisy RNNs and exact RNNs with action entropy regularization in the stable and volatile conditions. Noisy RNNs increase their inferred volatility in the volatile condition. **d**, Fraction of trials where noisy RNNs (left) and exact RNNs (right) choose the most rewarding lever at the beginning of the game (labelled action 1). Dashed lines indicate simulations of the best-fitting normative inference model, whereas unbroken lines indicate the behavior of RNNs. Only noisy RNNs approximate accurately the normative inference process.

As observed in humans, we found that noisy RNNs increased their inferred volatility in the volatile condition – an adaptive effect absent in exact RNNs (Fig. 17c, left). This effect was selective, in the sense that noisy RNNs increased much more their inferred volatility (by 310% on average) than their inferred trap probability (by 39%) in the volatile condition (Fig. 17c, middle). Finally, both types of networks showed increased choice stochasticity in the volatile condition (Fig. 17c, right). Comparing simulations of the best-fitting inference model to the behavior of noisy and exact RNNs confirmed that noisy RNNs approximate accurately the normative inference process, whereas exact RNNs showed significant deviations from the best-fitting process – due to their overly strong attractor states (Fig. 17d). These final results demonstrate that computation noise provides zero-shot, freestanding adaptation to volatility without any prior experience with volatile environments, nor any sophistication of the neural networks for dealing with such adverse conditions.

## Discussion

In cognitive psychology, general intelligence is not measured by the level of skill – or expertise – at any given task. Indeed, skill is known to depend strongly on prior experience, such that extensive training allows agents to reach extremely high levels of skill even when the acquired knowledge cannot be transferred to a different task – even a highly similar one. Recent definitions describe general intelligence as skill *acquisition* rather than skill itself^9,10^. In other words, intelligent agents are not necessarily the most skilled at any given task, but they are able to acquire new skills from previously acquired skills at low – to zero – cost. Artificial neural networks are prime examples of highly skilled agents with low generalization power. They perform extremely well on tasks with the same statistics to the ones on which they were trained, but they fare very poorly on conceptually similar tasks, or when confronted with adverse events in the tasks on which they were trained.

Increasing the generalization power of artificial agents constitutes one of the main challenges for AI research, and recent efforts have typically proceeded by engineering generalization abilities through purposeful sophistication of the artificial agents (e.g., hierarchical processes, meta-controllers…). Here we have explored an entirely different avenue for understanding generalization power in neural networks, by hypothesizing that this hallmark of general intelligence may be supported by an ubiquitous property of biological neural networks – namely computation noise – that is not shared with artificial neural networks. Across several tasks used to study higher cognition in humans and other animals (reasoning, learning, decision-making), we obtained converging evidence that computation noise boosts the generalization power of artificial neural networks by making them extremely resilient to a variety of adverse events and conditions entirely unseen during training, in a way that resembles human (and animal) cognition.

Our experimental approach for measuring the generalization power of our artificial agents consists in training them to perform a task A, and testing their ability to perform a more challenging task A* with adverse events absent from task A. Teaching an animal to perform a challenging task (whether it is reading or reversal learning) often requires to train the animal to excel at the different sub-components of the task. We thus reasoned that an intelligent agent trained on task A would be able to perform task A* with high accuracy without any prior experience – a form of ‘zero-shot’ acquisition of cognitive ability – by showing high cognitive resilience to adverse events specific to task A*. In this respect, artificial neural networks trained and tested with computation noise corrupting the update of their units were much more resilient – and thus much more intelligent – than their exact counterparts widely used in existing research.

The two classes of tasks we considered span two different areas of cognition research: the study of reasoning abilities during the manipulation of previously learned stimulus-response associations, and the study of online learning (‘meta-learning’) abilities during the active sampling of rewarded actions in novel environments. Therefore, during testing, our artificial agents became observers bound to integrate the presented cues in the weather prediction task, whereas they remained agents seeking rewards through interaction with their environment in the bandit task^32,38^. Despite these fundamental differences, we found that computation noise promotes very similar functions during reasoning and meta-learning – two cognitive abilities associated with substantial computation noise in humans^18–20^.

The structure of the computation noise we implemented in artificial neural networks was directly inspired by these recent observations in humans^18–20^. But instead of relying on an ‘algorithmic’ description of underlying computations as in earlier work (Bayesian inference in the weather prediction task, reinforcement learning in the bandit task), we adopted a neural network description which offers several advantages^29,30^. First, adding noise to optimal or near-optimal computations almost invariably degrades performance. For example, adding computation noise to the update of Bayesian beliefs triggers suboptimal behavior^18^, such that reducing computation noise – or suppressing it altogether – appears highly desirable from an evolutionary perspective when described in this global, process-level fashion. By contrast, here we added computation noise to the activity of each unit in a recurrent neural network. In such an interconnected information processing system, each unit does not compute alone any task variable, which is represented by populations of units. Therefore, computation noise added to individual units does not corrupt in an immediate, straightforward fashion the task variables computed by the neural network as a whole. As a consequence, the net effect of this local, unit-level computation noise on performance cannot be assumed to be detrimental. Our findings provide clear examples where computation noise promotes cognitive abilities in a zero-shot fashion – cognitive abilities that neural networks with exact computations lack.

These beneficial effects of computation noise on cognition call to reconsider the status of the ubiquitous internal variability of neural networks. Input noise (e.g., sensory noise) is widely seen as a hard constraint that neural networks have to cope with using efficient mechanisms^21^: ‘population’ coding to average out input noise^22^, or ‘efficient’ coding to limit its effects on the processing of natural stimuli^23,24^. We see computation noise as a biological constraint on neural activity. Indeed, in contrast to other stochastic heuristics designed to regularize the training of weights in deep neural networks (e.g., dropout^36^), computation noise was not artificially suppressed after training. It should therefore be seen as a constitutive property of neural networks, present during and after training, rather than as a engineered feature that can be turned off at will. Nevertheless, to test whether computation noise acts only as a performance-limiting constraint after training, we selectively suppressed computation noise during testing in ‘noise knocked-out’ agents, and compared their performance to the performance of their ‘wildtype’ (noisy) twins. We found that, in both reasoning and meta-learning agents, computation noise was necessary during testing to maximize cognitive resilience to adverse conditions. In other words, although we see computation noise as a genuine constraint on neural function, our findings suggest that general intelligence ‘rides’ on this constraint to promote high cognitive resilience when confronted with adverse, uncertain or volatile environments.

Others^42^ have theorized that suboptimal decisions are primarily due not to random noise (including computation noise), but to deterministic biases – i.e., systematic deviations from normative computations (e.g., Bayesian inference). This is not what we previously observed in humans^18,19^, and this is also not what we have observed here in noisy agents. In the weather prediction task, and as in humans, decomposing the suboptimality of noisy agents in terms of a ‘bias-variance’ trade-off revealed a large random variance term, and a small deterministic bias term. In other words, the behavior of noisy agents is truly noisy, and only weakly biased away from optimal computations. By contrast, the behavior of exact agents appeared highly biased in the same task: non-regularized agents decided on the basis of the first presented cue, whereas exact agents trained with action entropy regularization showed strong endogenous dynamics only moderately influenced by the presented cues. Furthermore, the fact that noisy agents incurred a small bias term is particularly noteworthy, since recurrent neural networks – even those with only 48 units, and thus about 3,000 weights to train – have weak inductive biases^43^. In other words, the same type of networks can learn a large variety of tasks, and exhibit a vast repertoire of possible dynamics and responses to the same input. From this perspective, it appears highly non-trivial that the many noisy agents we trained independently to learn probabilistic stimulus-response associations converged on highly similar behavior and activity patterns. Indeed, we observed much more inter-individual variability between exact agents trained to learn the same associations.

That noisy agents all converged on computations similar to Bayesian inference is even more compelling. Bayesian theories of brain function^44,45^ are prevalent in the recent literature, and they provide a unified framework for understanding animal behavior and neural computations across cognitive domains – such as multisensory integration^46^, motor control^47^, causal reasoning^48^ and language comprehension^49^. An intriguing possibility is that the ‘Bayesian brain’ is the natural consequence of noisy neural networks, where Bayesian computations correspond to a stable solution for a canonical task which constitutes the basis of several more complex tasks: the learning of rewarded stimulus-response associations.

Strikingly, in both reasoning and meta-learning tasks, the behavior of noisy agents was best fitted by Bayesian inference assuming non-zero volatility – i.e., the rate of changes in task statistics, a characteristic of adverse environments. The online monitoring of volatility exhibited by noisy metalearning agents is currently seen as the result of purposeful cognitive sophistication, through hierarchical inference in algorithmic models^37,41^, or the use of more complex units (e.g., long short-term memory units) in recurrent neural networks trained in volatile conditions^32^. In contrast to this view, our results show that computation noise promotes efficient adaptation to volatile environments in agents trained in perfectly stable conditions. Across the tasks we have tested, a fundamental property of computation noise appears to be the destabilization of attractor-like patterns of activity, an effect which can explain the higher performance of noisy agents across a range of adverse conditions without any explicit cognitive sophistication.

However, and despite its surprising benefits, we do not mean that computation noise is sufficient to explain the several complex forms of exploration observed in animals – humans in particular. The tasks we have considered here are laboratory experiments used to study learning and decision-making in controlled conditions. In contrast to exploration in the ‘wild’^50^, they require little to no exploration. Therefore, we do not argue that computation noise alone enables the efficient foraging of much more complex environments and replace, for example, directed exploration or curiosity-based modules^51–53^. Our claim focuses on the high cognitive resilience acquired by noisy agents in a zero-shot fashion – i.e., without any prior experience with adverse events during training.

Given that we implemented computation noise in artificial neural networks based on recent observations in humans^18,19^, it is particularly important to assess whether neural networks trained with computation noise show behavioral and neural patterns compatible with biological observations. A first important property of noisy agents is that the high cognitive resilience promoted by computation noise is robust to a large range of noise levels. From an engineering (AI) perspective, this property is convenient because it makes the setting of the noise level easier. But from a biological perspective, this property is crucial given the large inter-individual differences in cognitive parameters (including computation noise) observed across individuals. It would be extremely unlikely that computation noise promotes cognitive resilience if this effect was limited to very specific levels of noise. Furthermore, by studying the behavior of noisy artificial agents in tasks used to study higher cognition in animals, we could observe striking resemblances between noisy agents and human subjects. At the neural level, the activity patterns of noisy reasoning agents are highly consistent with neural observations from the lateral intraparietal cortex of macaque monkeys engaged in the weather prediction task^35^. Similarly, the neural dissociation between the representations of action outcomes and action values observed in noisy meta-learning agents is compatible with several neural observations from the prefrontal cortex^54^.

We have focused our study on the role of computation noise in learning and decision-making, by implementing it in a recurrent neural network used as a canonical model of associative cortical circuits in parietal and prefrontal regions^25–28^. However, several neural observations suggest that computation noise is present throughout the whole cortex, including sensory and motor regions. It is thus tempting to hypothesize, based on existing evidence, that introducing computation noise in deep feedforward convolutional models of the visual system^55^ or in multilayer perceptron models of sensorimotor circuits^56^ may promote the same sort of cognitive resilience to adverse conditions. Finally, exploring the benefits of computation noise for artificial intelligence systems appears like a promising, yet almost unexplored avenue in the search of artificial general intelligence^57,58^.

## Methods

### Neural network architecture

The artificial neural networks used for the weather prediction task and the bandit task are identical, and correspond to standard (Elman) recurrent neural networks^1^. Let us call *X*_*t*_ the input to the network, *Z*_*t*_ the recurrent state of the network and *Y*_*t*_ the output of the network. The recurrent neural network is governed by the following equations:

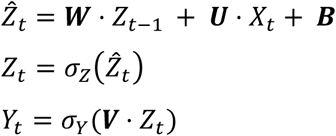

with *σ*_*Z*_ the hyperbolic tangent, *σ*_*Y*_ the softmax (sigmoid) function, and ⟨·⟩ the matrix multiplication operator. ***W, U, B*** and ***V*** are four matrices of network parameters adjusted during training. For the weather prediction task, *X*_*t*_ is a ‘one-hot’ vector encoding the presented cue (among 8 possible cues). For the bandit task, *X*_*t*_ is composed of the previous observed reward and a one-hot vector encoding the previous chosen action (among 2 possible actions). Following the first equation, the input *X*_*t*_ and the previous recurrent activity *Z*_*t*−1_ are integrated into an updated state 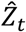 through matrix multiplication with weight matrices ***U*** and ***W*** (plus an additive bias term ***B***). This updated state 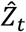 is then passed through a non-linearity *σ*_*Z*_ to give the updated recurrent activity *Z*_*t*_. This updated recurrent activity *Z*_*t*_ then projects to action probabilities *Y*_*t*_ through matrix multiplication with output weights ***V***, followed by the softmax *σ*_*Y*_ operator. For both tasks, we used *K* = 48 units in the recurrent layer, resulting in 2,832 free parameters to adjust during training in the weather prediction task, and 2,592 free parameters in the bandit task.

### Objective functions

In both tasks, the objective functions used for training the networks are derived from obtained rewards. In the weather prediction task, where all stage 1 trials are independent from one another, the objective function writes as follows:

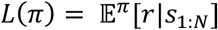

with *s*_1:*N*_ the *N* presented cues (*N* = 5), *r* the obtained reward, and *π* the ‘action policy’ giving the probability of each action (i.e., the output layer of the neural network). In the bandit task where the successive trials are dependent, the objective function writes as follows:

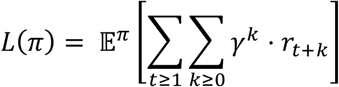

with *t* the trial number, *γ* ∈ [0,1] the discount factor, *r*_*t*+*k*_ the obtained reward at trial *t* + *k* (with *k* a positive integer) and *π* the action policy used by the neural network. Having no prior assumptions regarding *γ*, we used *γ* = 0.5.

### Training procedure

The recurrent neural networks have a set of parameters (the matrices ***U, V, W*** and ***B*** described in the **Model architecture** subsection) that we trained using the REINFORCE algorithm^2^. This training procedure relies on a direct differentiation of the objective functions. In the weather prediction task, the gradient of the objective function writes as follows:

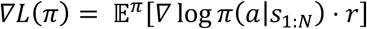

with *s*_1:*N*_ the presented cues, *r* the obtained reward, *π* the action policy of the neural network and *a* the chosen action. In the bandit task, the gradient of the objective function writes as follows:

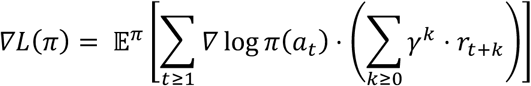

with *t* the trial number, *a*_*t*_ the chosen action at trial *t, r*_*t*+*k*_ the observed reward at time *t* + *k* (*k* a positive integer), *π* the action policy of the neural network and *γ* the discount factor.

On both tasks, we trained 50 artificial agents using the same stochastic training procedure. The behavior and activity patterns of the 50 trained agents were entered as repeated measures in all analyses reported below. The stochastic gradient descent (SGD) procedure was performed with the RMSProp optimizer^3^ and a learning rate of 0.0001. We set the total number of training steps to 50,000 for the weather prediction task, with each step consisting of 100 independent trials. We also set it to 50,000 for the bandit task, with each step consisting of one game of 100 trials. Asymptotic performance was reached at the end of the SGD procedure in both cases.

### Introducing computation noise in neural networks

We introduced in this study three types of recurrent neural networks (RNNs). The two first types feature exact (noise-free) computations: in other words, their dynamics are governed by exact applications of the equations described in the **Model architecture** subsection. The first type, referred to as ‘no regularization’, is defined by the dynamics and objective functions described above.

The second type, referred to as “action entropy regularization”, is identical to the “no regularization” type in its architecture but is trained using an objective function that encourages explicitly action policies with high entropy (random exploration). This is done by adding the entropy of the action policy *π* as an additive term to the objective function *L*(*π*):

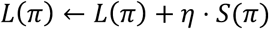

with *S*(·) the entropy function and *η* a positive scaling factor. This form of action entropy regularization is commonly used for training deep reinforcement learning networks^4,5^.

The third type, referred to as “computation noise”, is identical to the “no regularization” type in its objective function but features random noise in the equations that govern its dynamics. We implemented computation noise in the network dynamics by updating the activity of each unit in the network in an imprecise fashion. Let *Z*_*t*−1_ be the recurrent activity of the network at time *t* − 1 and 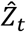 the updated state at time *t* before the non-linearity *σ*_*Z*_. For the weather prediction task, based on recent behavioral observations of computation noise in humans, we corrupted the updated state of each unit in the network with i.i.d. normally distributed noise. More precisely, we sampled the noisy updated state 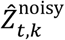 from a normal distribution of mean equal to the result of the exact (noise-free) update 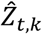, and of standard deviation *σ*:

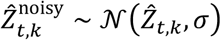

with 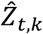 the activity of unit *k* (*k* ≤ *K*, with *K* = 48 the total number of units in the network) such that 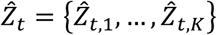. For the bandit task, based also on recent behavioral observations in humans, we assumed that the standard deviation *σ* of computation noise scales with the magnitude of update at each unit – i.e., following Weber’s law:

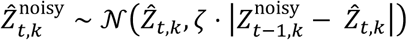

with *ζ* > 0 the scaling coefficient of computation noise also referred to as its ‘Weber fraction’. Computation noise thus has slightly different structures in the two tasks: a constant-scaling structure in the weather prediction task where noisy agents manipulate previously learnt stimulus-response associations, and a multiplicative-scaling structure in the bandit task where noisy agents track rewarded actions through interaction with their environment. Constant-scaling noise in the weather prediction task could reflect, at least in part, task-irrelevant input which effectively corrupts the computation of task-relevant variables. On the other hand, the multiplicative-scaling noise implemented in the bandit, ‘meta-learning’ task reflects stochastic trial-by-trial reconfigurations of the network following each outcome. Once the noisy updated state 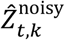 is sampled, the non-linear activation function *σ*_*Z*_ is applied exactly as in exact neural networks. Critically, we implemented computation noise as a constraint in the network and thus made the SGD procedure unaware of the source of this noise, by nullifying the gradients that arise from its associated variance.

### Experimental procedures for the weather prediction task

The training task (stage 1) is composed of independent trials of *n* = 5 samples of the same cue (sampled uniformly among the 8 cues shown in Fig. 2). On each trial, the agent is presented with one of the cues for 5 samples, after which the agent has to choose between two actions. The agent receives a positive (+1) or negative (−1) reward as a function of the probabilistic association between the presented cue and the chosen action. In stage 1, optimal behavior consists in choosing the ‘greedy’ action that maximizes the probability of obtaining a positive reward.

Once the network is trained through SGD (as described in the **Training procedure** subsection), we fix its weights and test it in the weather prediction condition (stage 2). On each trial, the agent is now presented with sequences of different cues on each sample, with *n* = [2,…,16] samples. After each sequence, the agent has to choose between the same two actions the one most likely to be associated with reward given the presented cues. In this second stage, each cue taken in isolation is associated with the same reward probabilities as in the first stage, but reward probabilities have to be combined across presented cues to identify the best action associated with the sequence of cues as a whole. The difficulty of this ‘weather prediction’ stage can be manipulated by varying the fraction of conflicting cues (i.e., cues associated in stage 1 with a different rewarded action than the sequence as a whole).

In practice, to determine which cues would be presented on a particular trial, we defined two categories, each associated with one of the two actions being rewarded at the end of the trial. At the beginning of the trial, one of the two categories was randomly selected for subsequent sampling. The distributions of cues associated with each category were defined such that the conditional probability that a cue is sampled matches its reward probability experienced in stage 1. Additionally, in some instances of the weather prediction task, we introduced a reversal in the category being sampled after 8 samples, such that the rewarded action at the end of the trial is inconsistent with the cues presented during the first 8 samples (before the reversal), but consistent with the cues presented during the last 8 samples (after the reversal).

### Experimental procedures for the bandit task

The training task consists of a two-armed bandit game. On each trial of a game, the agent is presented with a slot machine with two levers to choose from. The agent receives a positive (+1) or negative (−1) reward as a function of the reward probability associated with the chosen lever in the current game (0.95 for the best lever, 0.05 for the worst lever), which remains fixed over the course of the game. The most rewarded lever is reset randomly at the beginning of each game, such that the agent needs to learn which lever is most rewarded in every single game.

Once the network is trained through SGD (as described in the **Training procedure** subsection), we fix its weights and test it in several variants of the canonical bandit task described above. In the first variant (early trap task), we introduced a number of misleading outcomes, or ‘traps’, at the beginning of each game. In the second variant (reversal learning task), we introduced a reversal in the reward probabilities associated with the two levers after 25 trials. In the third variant (restless bandit task), the reward probabilities associated with the two levers drift randomly over the course of each game. And in the fourth and last variant, we introduced a ‘volatile’ condition in which the reward probabilities associated with the two levers switch every 25 trials.

### Fitting normative behavior in the volatile weather prediction task

We fitted a model derived from the normative Bayesian inference process assuming non-zero volatility^6^ (i.e., possible reversals in the rewarded action in the middle of a sequence of cues) to the behavior of noisy agents. The model has two free parameters *θ*: a volatility parameter *v* which corresponds to the reversal probability assumed by the model, and a decision gain *β* which corresponds to the inverse temperature of the softmax action policy used by the model. Briefly, beliefs are defined in this model as the log-posterior odds ratio between the possible sources of cues given all trials until the current trial *t*. Let us call *L*_*t*−1_ the log-posterior odds ratio at trial *t* − 1. Given the cues presented at trial *t*, which together define a log-likelihood odds ratio LLR_*t*_, the log-posterior odds ratio is updated through the following equation:

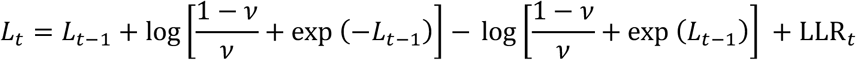

with *v* the volatility parameter. Choice is then made at the end of trial *t* based on the log-posterior odds ratio *L*_*t*_ through a softmax action policy with inverse temperature *β*. The two free parameters *v* and *β* were fitted to the behavior of noisy agents through maximum likelihood estimation (MLE) separately for each artificial agent.

### Fitting normative behavior in the volatile bandit task

We fitted a model derived from the normative Bayesian inference process assuming non-zero volatility^7^ to the behavior of noisy and exact agents. The model has three free parameters *θ*: the volatility *v* assumed by the model, the trap probability *ρ* assumed by the model, and the decision gain *β* used by the softmax action policy of the model. Briefly, the model tracks the hidden state *z*_*t*_ characterizing the binary action-outcome state of the task at the current trial *t*: *z*_*t*_ = 1 if action 1 is the highest rewarded one, and *z*_*t*_ = 0 otherwise. Based on the prior belief on the hidden state *z*_*t*_, *p*(*z*_*t*_ | *r*_1:(*t*−1)_, *a*_1:(*t*−1)_, *θ*), the action *a*_*t*_ is chosen by applying a softmax policy with inverse temperature *β* to the prior belief. This chosen action *a*_*t*_ leads to an obtained reward *r*_*t*_. The prior belief from trial *t* is then updated to a posterior belief by incorporating the likelihood of the obtained reward:

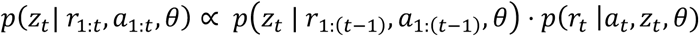

with 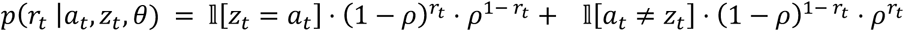 the likelihood function. The posterior belief at trial *t* is then updated to a prior belief at trial *t* + 1:

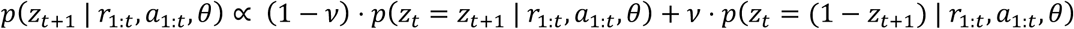

From this prior belief at trial *t* + 1, the inference process is applied iteratively. The three free parameters *v, ρ* and *β* were fitted to the behavior of noisy agents through maximum likelihood estimation (MLE) separately for each artificial agent.

## Author contributions

C.F. conceptualized the study and developed the neural networks. C.F. and V.W. designed and performed the experiments. C.F. and V.W. conducted the data analyses. C.F. and V.W. interpreted the results and wrote the manuscript. V.W. acquired funding.

## Acknowledgments

This work was supported by a starting grant from the European Research Council awarded to V.W. (ERC-StG-759341), and a collective grant from the Agence Nationale de la Recherche awarded to the Département d’Études Cognitives (ANR-17-EURE-0017).

