## Supplementary Information for "Computation noise promotes cognitive resilience to adverse conditions during decision-making"

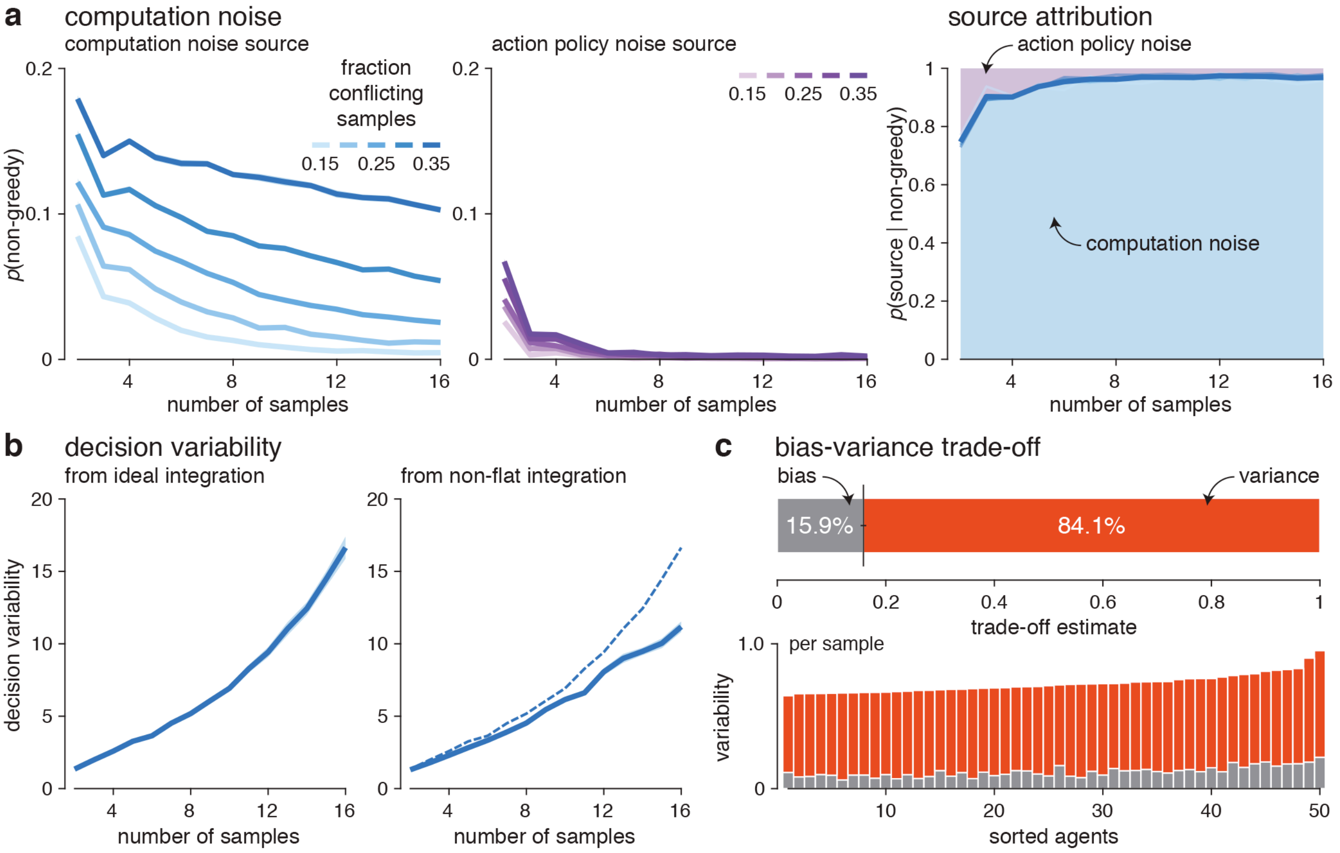


**Supplementary Fig. 1 | Characterization of behavioral variability in the weather prediction task. a**, Left: fraction of non-greedy decisions triggered by the computation noise source (left) and action policy noise source (right) of noisy RNNs ($\sigma$ = 2), for increasing numbers of samples in the trial. Non-greedy decisions are defined as decisions where the agent does not choose the action predicted by applying exact (noise-free) updates of recurrent activity (triggered by computation noise), or decisions where the agent does not choose the action predicted by a purely deterministic (argmax) action policy (triggered by action policy noise). Non-greedy decisions triggered by both noise sources decrease with sequence length, and increase with the fraction of conflicting samples in the sequence. Right: relative contributions of computation noise (blue-shaded area) and action policy noise (violet-shaded area) to non-greedy decisions. As in humans, most non-greedy decisions are triggered by computation noise. **b**, Decision variability as a function of the number of samples. Decision variability is defined as the estimated variance of deviations between the decision variable used by the agent and the ideal log-posterior computed by a Bayesian decision-maker. Left: decision variability from ideal (flat) integration. Right: decision variability from the empirical (non-flat) integration kernel estimated for each agent. In both cases, decision variability grows parametrically with the number of samples. **c**, Bias-variance trade-off of the choice suboptimality of noisy RNNs. Top: mean bias-variance trade-off across 50 noisy RNNs, estimated by comparing the actions taken by each noisy RNN to two repetitions of the same sequence of cues. The bias terms reflects systematic deviations from Bayesian inference, whereas the variance term reflects random deviations due to computation noise. As in humans, the accuracy of noisy RNNs is bounded by the variance term (84.1% of the overall suboptimality). Bottom: separate bias (gray) and variance (red) terms across 50 noisy RNNs. All agents exhibit similar bias and variance terms.

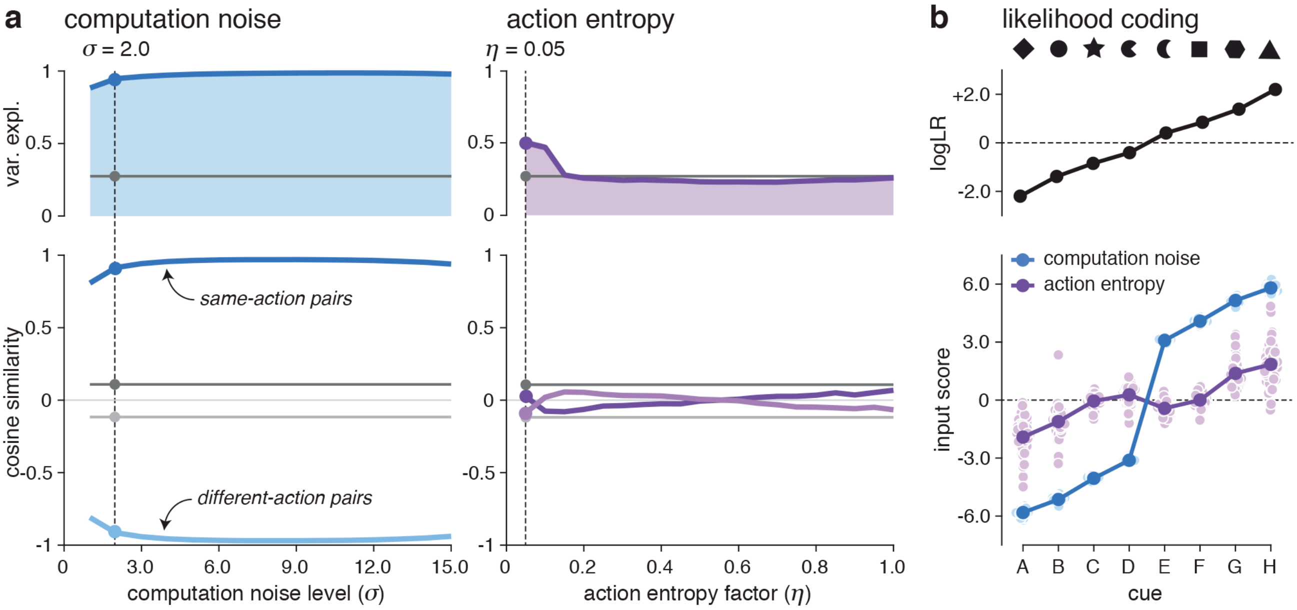


**Supplementary Fig. 2 | Line-based representation of input in the weather prediction task. a**, Principal component analysis of input weights in noisy RNNs (left) and exact RNNs with action entropy regularization (right). Top: overall variance explained by the first PC of input weights across computation noise levels (left) and action entropy factors (right). The first PC captures more than 90% of the overall variance of input weights for noisy RNNs, but only 25% for exact RNNs with action entropy regularization and for non-regularized RNNs (in gray). Bottom: cosine similarity between the input weights of cue pairs associated with the same action (same-action pairs), and cue pairs associated with different actions (different-action pairs), across computation noise levels (left) and action entropy factors (right). In noisy RNNs, cues associated with the same action project in the same direction and cues associated with different actions project in opposite directions. **b**, Encoding of the log-likelihood ratio provided by each cue by input weights. Top: ideal log-likelihood ratio (logLR) provided by each cue. Bottom: input score associated with each cue. The input score is defined as the projection of the input weights associated with each cue on the first PC. Input scores revealed an ordering of the cues as a function of their log-likelihood ratios in noisy RNNs (blue), not in exact RNNs (violet).

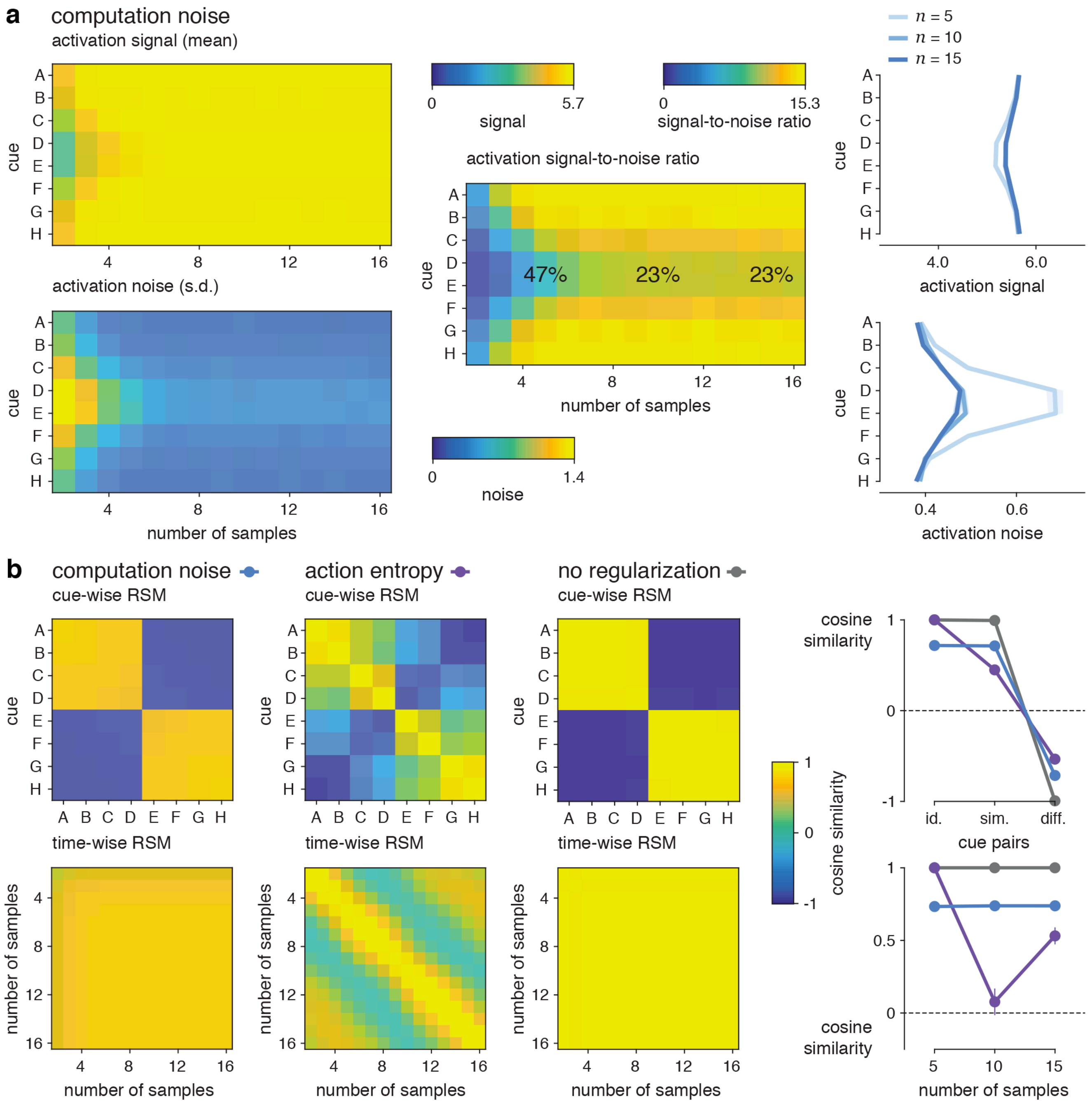


**Supplementary Fig. 3 | Characterization of activity patterns in the weather prediction task. a**, Statistics of activity patterns in noisy RNNs ($\sigma$ = 2) during stage 1 (single cue presentation) for increasing numbers of samples. Activation is defined as the projection of cue-wise activity patterns on their mean directions. Top: activation signal (mean) for each cue (y-axis). Activation signal saturates after 5 samples and does not distinguish between cues of different reliabilities. Bottom: activation noise (standard deviation) for each cue. Activation noise decreases with the number of presented samples and increases in response to less reliable cues for any presentation time. Middle: activation signal-to-noise ratio (SNR). The SNR decreases in response to less reliable cues for any presentation time (by 47% for 5 samples, by 23% for 10 samples and more). Right: profiles of activation signal (top) and noise (bottom) across cues for 5, 10 and 15 samples. Activation noise varies substantially more than activation signal as a function of cue reliability. **b**, Representational similarity analysis (RSA) during stage 1. Top: cue-wise RSA (similarity between activity patterns triggered by different cues) for noisy RNNs (left; $\sigma$ = 2), exact RNNs with action entropy regularization (middle; $\eta$ = 0.05) and non-regularized RNNs (right). Bottom: time-wise RSA (similarity between activity patterns triggered by the same cue between different time samples) for the same three types of networks. Exact RNNs trained with action entropy regularization do not show a stable representation of each cue over time.

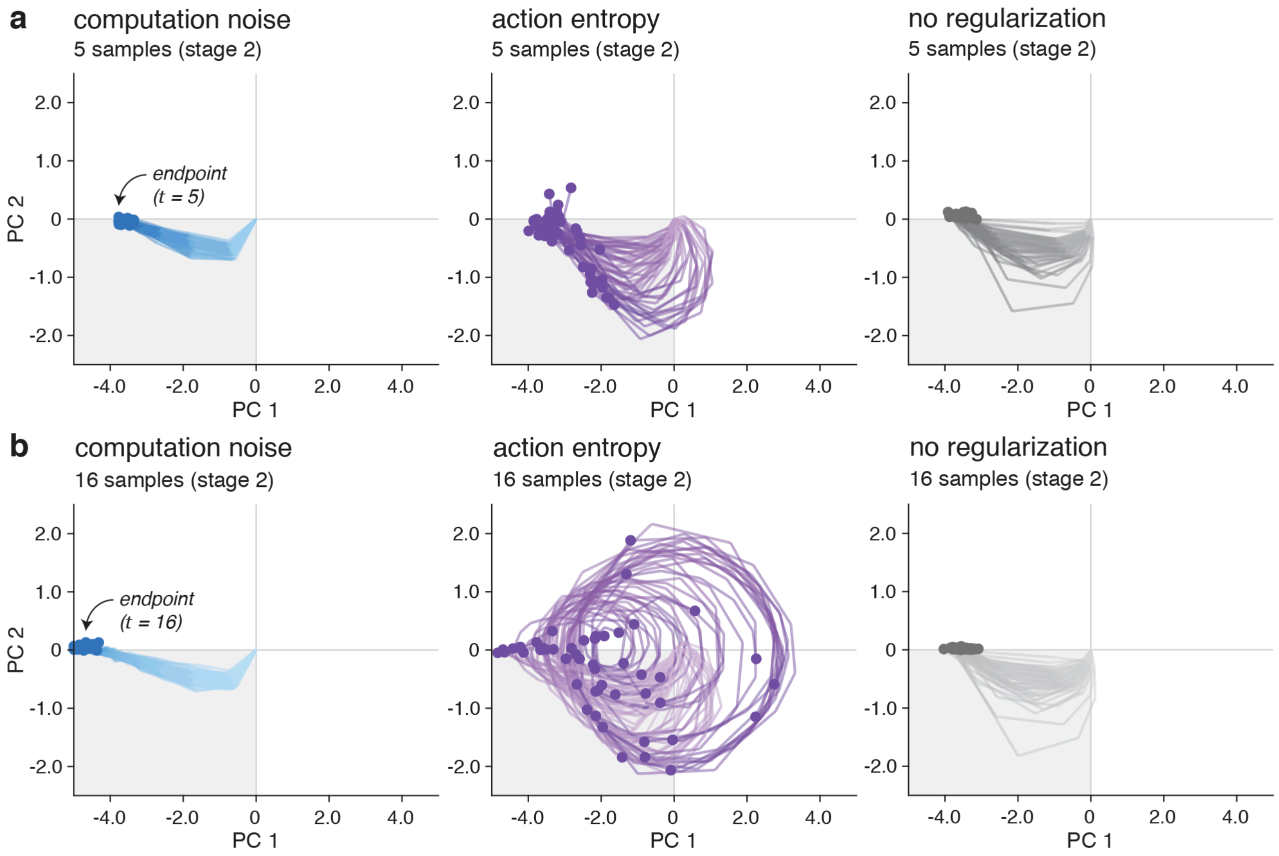


**Supplementary Fig. 4 | Trajectories of activity patterns in the weather prediction task.** Activity trajectories of individual RNNs in the two-dimensional space defined by their first principal components. The trajectories displayed here are obtained during stage 2 (weather prediction) for 1,000 trials (identical trials across the 50 RNNs of each type). PC 1 activity (x-axis) is signed such that it correlates positively with the log-posterior after $n$ = 5 samples (the presentation time used for training in stage 1). PC 2 activity (y-axis) is signed such that it correlates positively with the log-likelihood of presented cues. **a**, Activity trajectories in response to sequences of $n$ = 5 samples (the number used for training the network weights), for noisy RNNs (left; $\sigma$ = 2), exact RNNs with action entropy regularization (middle; $\eta$ = 0.05) and non-regularized RNNs (right). **b**, Activity trajectories in response to sequences of $n$ = 16 samples, for the same three types of networks. The activity of noisy RNNs converges rapidly toward highly consistent endpoints across trained agents, endpoints whose distance from zero scales with the number of presented samples. The activity of exact RNNs trained with action entropy regularization shows wild swings across the activity space, and the activity of non-regularized RNNs converges toward fixed endpoints independent of the number of presented samples.

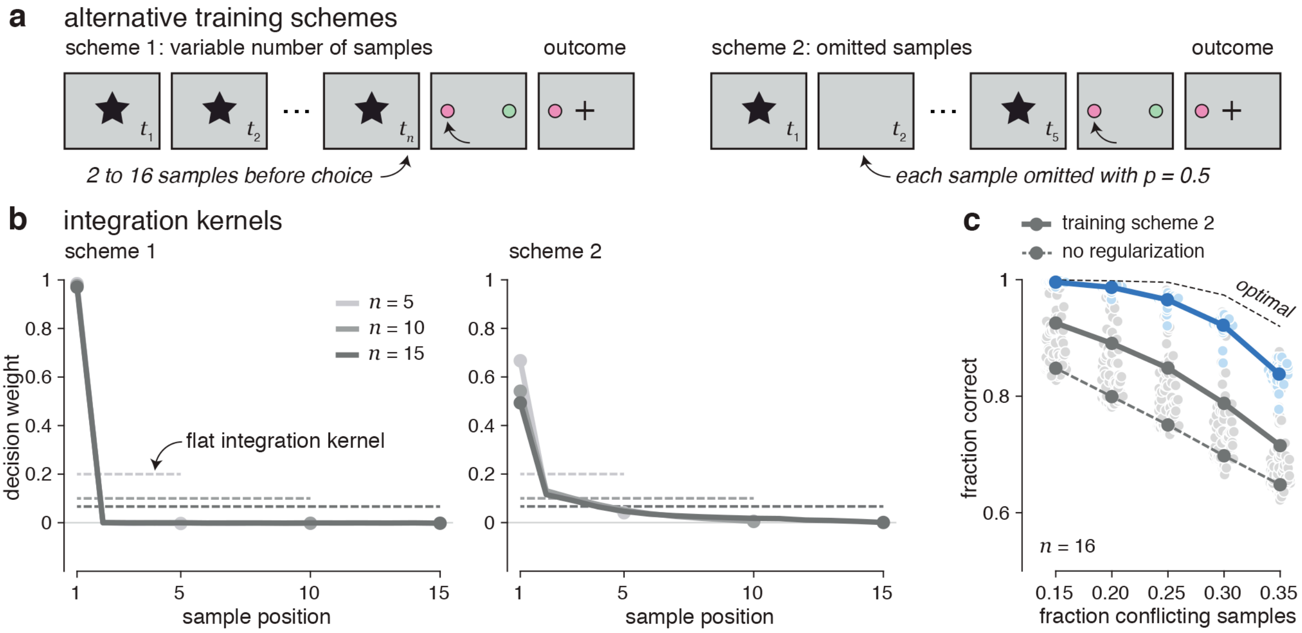


**Supplementary Fig. 5 | Alternative training schemes for exact RNNs. a**, Description of alternative training schemes in stage 1. Left: training scheme 1, using a variable number of samples ($n$ = 2 to 16 samples) instead of a fixed number of samples ($n$ = 5). Right: training scheme 2, using a probabilistic omission of cue presentation at each sample with probability $p$ = 0.5. **b**, Psychophysical integration kernels of non-regularized RNNs trained using alternative scheme 1 (left) and alternative scheme 2 (right). Training scheme 1 does not improve the extreme primacy bias shown by non-regularized RNNs, whereas training scheme 2 reduces the severity of the bias. **c**, Fraction of correct decisions in stage 2 ($n$ = 16 samples) as a function of the fraction of conflicting samples for noisy RNNs (blue curve) and non-regularized RNNs trained either using the same scheme as noisy RNNs (dashed curve) or using the alternative scheme 2 (unbroken curve). Non-regularized RNNs trained using this alternative scheme remain substantially worse than noisy RNNs in stage 2.

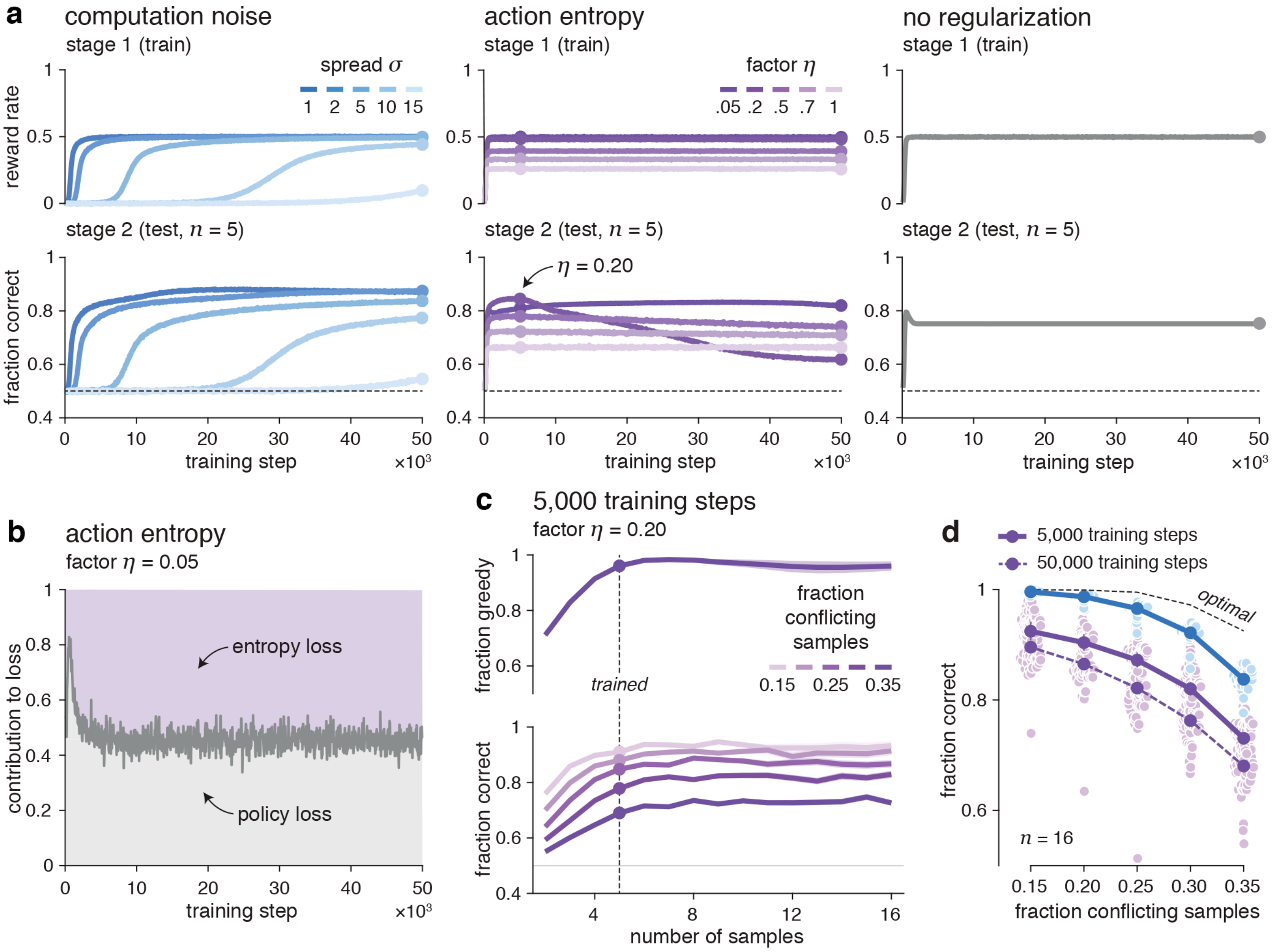


**Supplementary Fig. 6 | Performance in the weather prediction task throughout training. a**, Performance in stage 1 (reward rate used for training the network weights) and stage 2 (fraction of correct decisions for sequences of 5 samples with 25% of conflicting samples) as a function of training step, for noisy RNNs (left), exact RNNs with action entropy regularization (middle) and non-regularized RNNs (right). Noisy RNNs show synchronous improvements in stage 1 (training condition) and stage 2 (testing condition) during training of the network weights: a clear testimony of 'zero-shot' learning of the weather prediction task. By contrast, the performance of exact RNNs trained with action entropy regularization shows in stage 2 an early rise followed by a progressive decline characteristic of overfitting in stage 1. This late decline is particularly pronounced for exact RNNs with $\eta$ = 0.20, whose performance peaks after about 5,000 training steps. Non-regularized RNNs converge rapidly on an extreme primacy bias. **b**, Relative contributions of the action policy term (gray area) and the entropy term (violet area) to the overall loss for exact RNNs trained with $\eta$ = 0.05, whose performance in stage 2 does not decline throughout training. The policy loss outweighs the entropy loss at the beginning of training, and the two loss terms converge toward equal contributions after 5,000 training steps. **c**, Fraction of greedy decisions in stage 1 (top) and fraction of correct decisions in stage 2 (bottom) as a function of the number of samples for exact RNNs trained with $\eta$ = 0.20 for 5,000 training steps (instead of 50,000). The early stopping of training before overfitting does not yield hallmarks of Bayesian inference in exact RNNs. Accuracy in stage 2 saturates rapidly for more than 5 presented samples. **d**, Fraction of correct decisions in stage 2 ($n$ = 16 samples) as a function of the fraction of conflicting samples for noisy RNNs (blue curve) and exact RNNs training with $\eta$ = 0.20 for 50,000 training steps (dashed curve) and 5,000 training steps (unbroken curve). Non-overfitted exact RNNs remain substantially worse than noisy RNNs in stage 2.

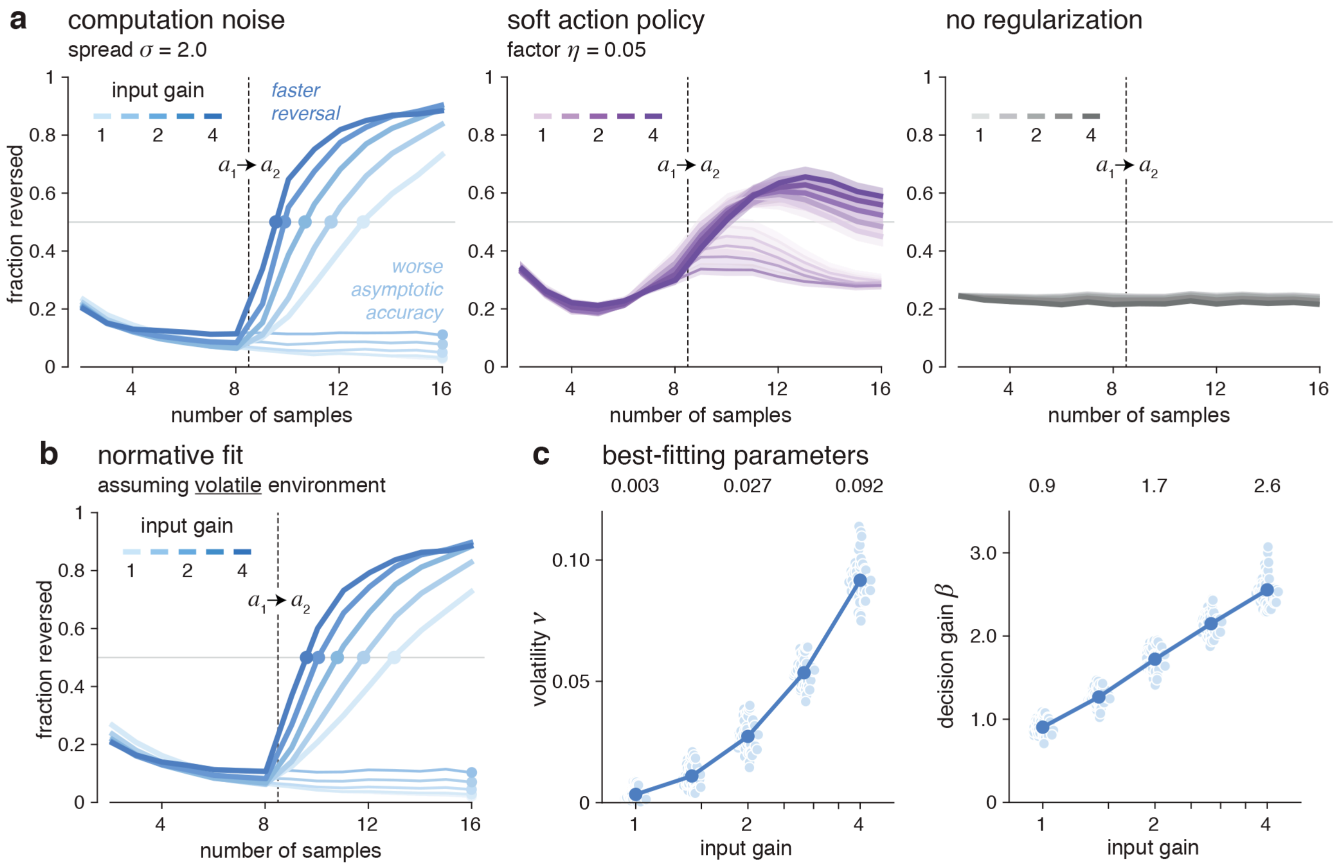


**Supplementary Fig. 7 | Training-free control of inferred volatility by input gain. a**, Fraction of trials where action 2 (the rewarded action for sequences longer than 8 trials) is chosen by noisy RNNs (left; $\sigma$ = 2), exact RNNs with action entropy regularization (middle; $\eta$ = 0.05) and non-regularized RNNs (right) in condition 1 (thin lines; no reversal) and condition 2 (thick lines; reversal after 8 samples) of the volatile weather prediction task, for varying levels of input gain (baseline = 1). Noisy RNNs reverse their behavior faster with increasing input gain, at the expense of worse asymptotic accuracy in sequences without reversal. **b**, Fits of normative integration assuming non-zero volatility to the behavior of noisy RNNs driven by increasing input gain. The faster adaptation of noisy RNNs to mid-sequence reversals with increased input gain is well captured by the normative inference process assuming non-zero volatility. **c**, Best-fitting estimates of inferred volatility (left) and decision gain (right) for noisy RNNs driven by varying levels of input gain. An increase in input gain was captured by a whopping 30-fold increase in the volatility assumed by the model.

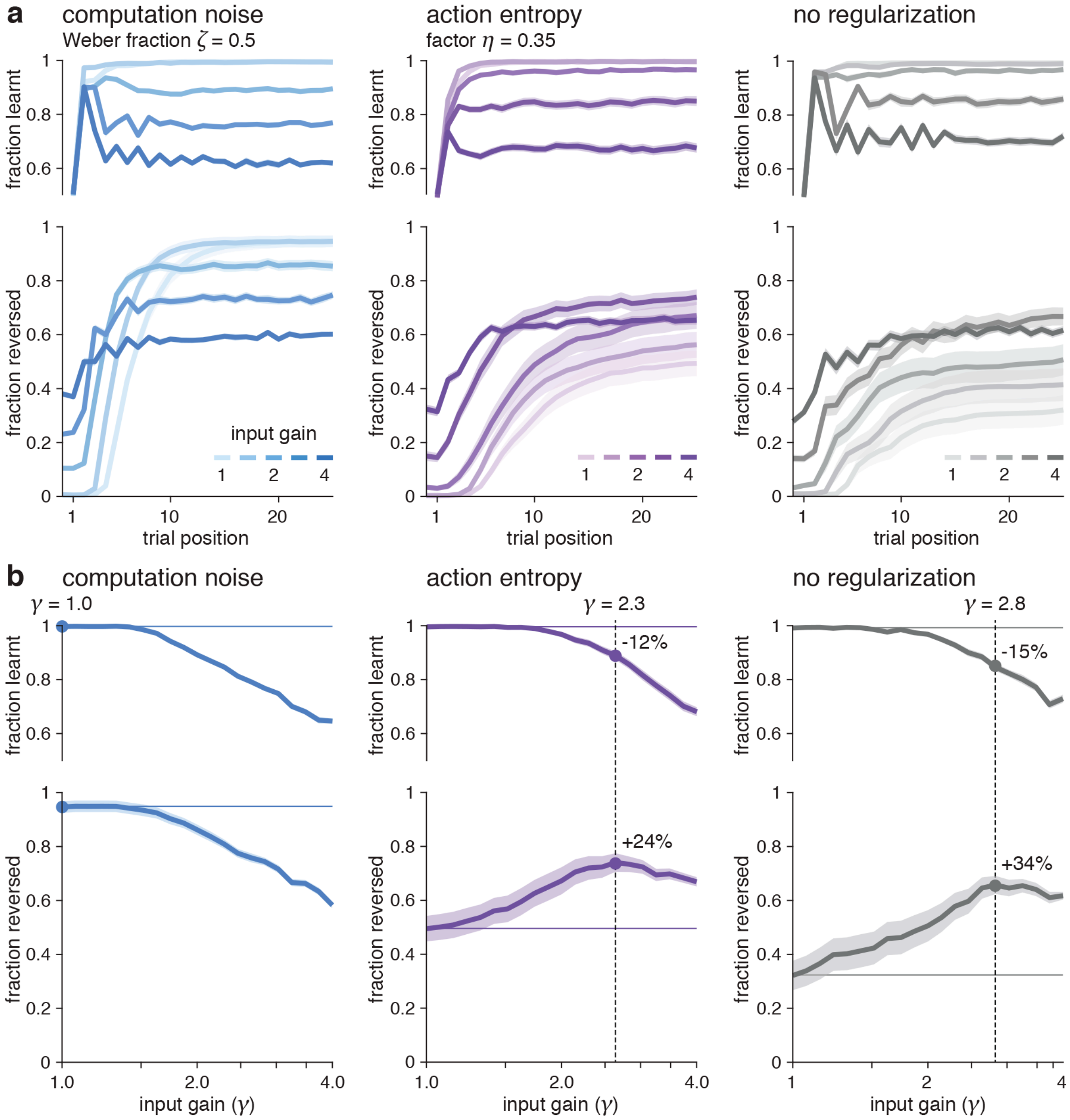


**Supplementary Fig. 8 | Training-free control of learning rate by input gain.** Manipulation of input gain in meta-learning agents trained in a bandit task with fixed reward probabilities 0.95 and 0.05 and tested in the reversal bandit task. **a**, Learning curves (before reversal, top) and reversal curves (after reversal, bottom) for noisy RNNs (left; $\zeta$ = 0.5), exact RNNs with action entropy regularization (middle; $\eta$ = 0.35) and non-regularized RNNs (right) driven by varying levels of input gain. Increasing the input gain to the network increases the effective learning rate of all types of networks: increased reversal learning performance for moderate increases in input gain (faster reversals for noisy RNNs, fewer missed reversals for exact RNNs), traded against decreased performance during initial learning. Increasing input gain further yields agents with poor performance in both stages of the reversal learning task. **b**, Asymptotic learning rate (before reversal, top) and asymptotic reversal rate (after reversal, bottom) as a function of input gain for noisy RNNs (left), exact RNNs with action entropy regularization (middle) and non-regularized RNNs (right). Increasing input gain beyond $\gamma \approx 1.5$ degrades learning and reversal performance in noisy RNNs, but increases reversal performance in exact RNNs up to $\gamma \approx 2.8$, at the expense of worse learning performance.

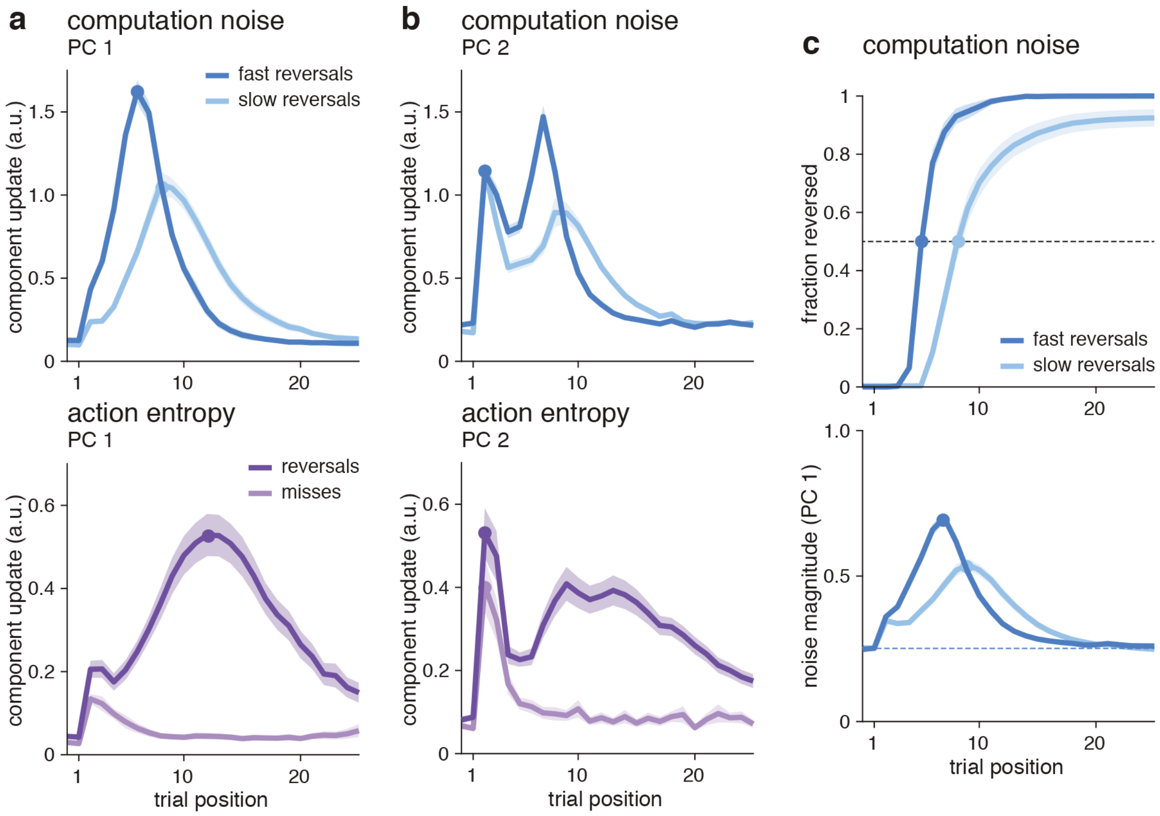


**Supplementary Fig. 9 | Relation between reversal dynamics and behavior. a**, Updates of PC 1 activity in games associated with fast vs. slow behavioral reversals in noisy RNNs (top; $\zeta$ = 0.5), and games associated with detected vs. missed reversals in exact RNNs with action entropy regularization (bottom; $\eta$ = 0.35). Trial 1 corresponds to the first trial after the reversal in reward probabilities. In noisy RNNs, PC 1 activity shows larger updates in games associated with fast behavioral reversals. In exact RNNs, PC 1 activity shows almost no update in games associated with missed reversals, but large updates in games associated with detected reversals. **b**, Updates of PC 2 activity in games associated with fast vs. slow reversals in noisy RNNs (top), and games associated with detected vs. missed reversals in exact RNNs with action entropy regularization (bottom). In noisy RNNs, the early update of PC 2 in the first trial following a reversed outcome does not distinguish between fast and slow behavioral reversals. In exact RNNs, the early update of PC 2 predicts whether the network subsequently detects or misses the reversal. **c**, Time courses of choice behavior (fraction reversed, top) and noise magnitude (bottom, defined as the unsigned difference between noisy and noise-free realizations of the update of PC 1 activity at trial $t$) in games associated with fast vs. slow reversals in noisy RNNs. Fast behavioral reversals are preceded by larger noise magnitude on PC 1 activity.
